# Whole body MondoA deletion protects against diet-induced obesity through uncontrolled multi-organ substrate utilization and futile cycling

**DOI:** 10.1101/2025.10.06.680559

**Authors:** Justin H. Berger, Allison N. Lau, Larry C. James, Renee Taing, Byungyong Ahn, Xiaofei Yin, Tomoya Sakamoto, Kirill Batmanov, Olivia Jordan, Jiten Patel, Jeffrey Zhou, Paul M. Titchenell, Brian N. Finck, Gregory J. Tesz, Daniel P. Kelly

## Abstract

**Objective:** Delineating the nodal control points that maintain whole-body energy homeostasis is critical for understanding potential treatments of obesity and cardiometabolic diseases. The nutrient-sensing transcription factor MondoA is a regulator of skeletal muscle fuel storage, where muscle-specific inhibition improves glucose tolerance and insulin sensitivity. However, the role of MondoA in whole body energy metabolic homeostasis is not understood.

**Methods:** Generalized MondoA knockout (gKO) mice were generated and assessed for glucose tolerance and insulin sensitivity, body composition, energy expenditure, cold tolerance, and tissue specific transcriptional changes in response to high fat diet. Complementary studies in cultured human adipocytes assessed the impact of MondoA deficiency on substrate utilization and lipolysis.

**Results:** gKO mice are protected from diet-induced obesity and insulin resistance, through increased whole body energy expenditure. gKO mice exhibit reduced brown and inguinal white adipose tissue mass, without evidence of beiging. The gKO mice are hyperlactatemic and isolated MondoA-deficient adipocytes have increased 2-deoxyglucose uptake and glycolytic function. Lastly, gKO mice and KO adipocytes display increased circulating glycerol relative to free fatty acids in response to adrenergic stimulus consistent with elevated re-esterification. However, this phenotype is not recapitulated in adipocyte-specific KO mice.

**Conclusions:** MondoA deficiency alters cellular sensing of nutrient availability and storage/utilization mechanisms. In the whole-body setting, this results in increased energy expenditure, potentially related to increased glucose uptake and glycolytic flux driving glycerol synthesis to supply high rates of lipolysis and lipid re-esterification. These results suggest that MondoA functions to maintain fuel storage and when lost, inter-organ futile cycling ensues.

Graphical Abstract.
1) Global MondoA deficiency drives 2) tissue glucose uptake which in skeletal muscle is 3) converted and excreted as lactate, while in adipose tissue 4) triglyceride re-esterification requires 5) de novo glycerol synthesis to feed into the futile cycle.

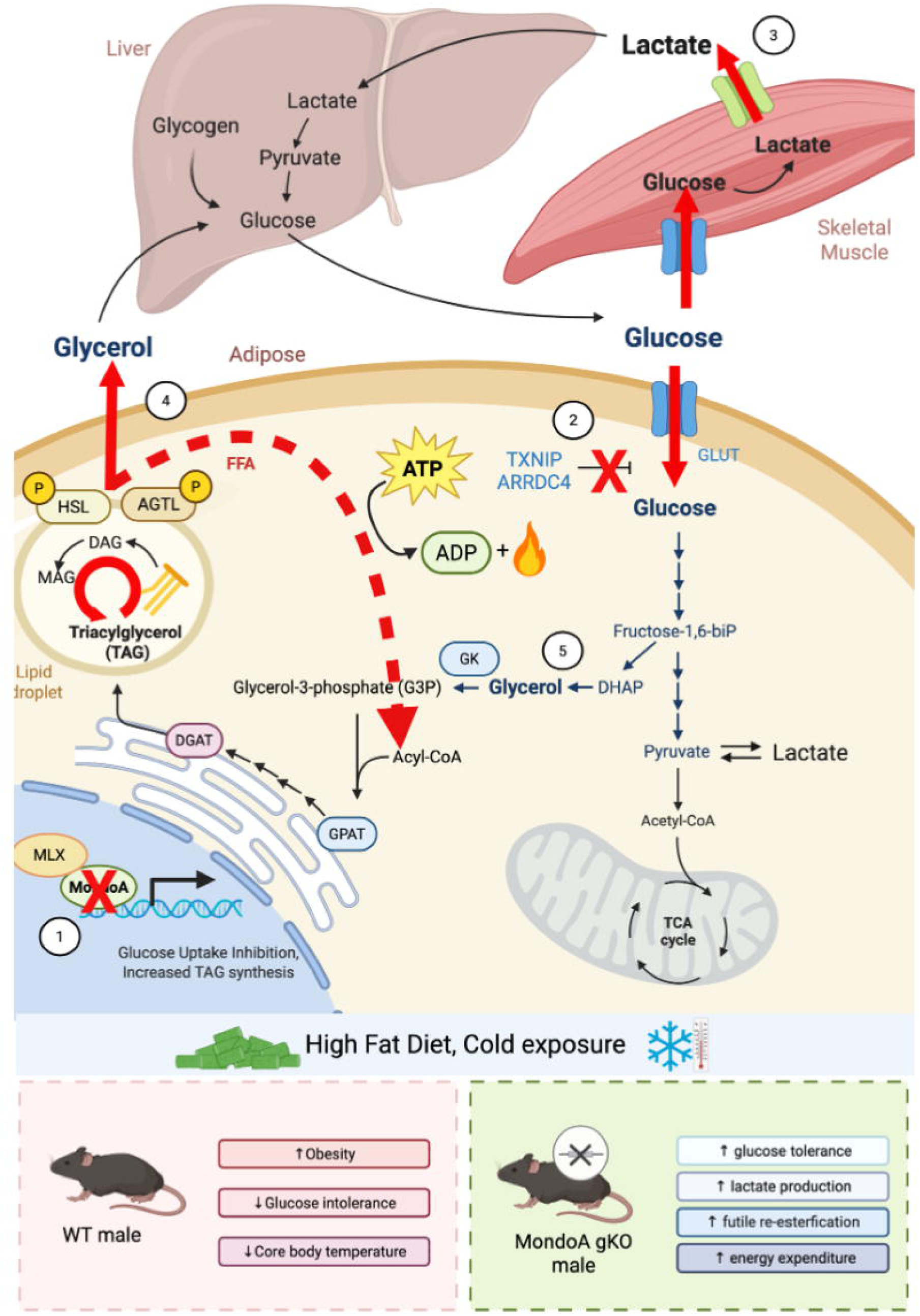

## Introduction

Metabolic dysregulation occurs in a broad array of human chronic diseases, including obesity and diabetes, but also nutrient-wasting states such as cancer, heart failure and age-related sarcopenia. The regulation of organismal energy homeostasis requires multiple complex layers of feedback mechanisms that involve nutrient sensing and cross-tissue signaling to determine rates of whole-body energy expenditure, which partially explains why therapeutic targeting of energy expenditure in obesity and cachectic states has been challenging.

Previously, we demonstrated that the transcription factor MondoA, a skeletal muscle-enriched paralog of carbohydrate-response element binding protein (ChREBP), serves as a nutrient-sensing regulator of skeletal muscle glucose uptake and fuel storage. Specifically, muscle-specific genetic targeting of MondoA resulted in enhanced glucose uptake and fatty acid utilization, and reduced high fat diet-induced skeletal muscle lipid accumulation due to deactivation of MondoA targets *Txnip* and *Arrdc4*. The muscle-specific MondoA-deficient mice were more glucose tolerant and insulin sensitive on a high fat diet compared to wild-type counterparts, without a difference in weight gain. In addition, short-term administration of a small molecule inhibitor of MondoA phenocopied this muscle phenotype. suggesting that MondoA or downstream targets could prove to be new therapeutic targets [1, 2]. MondoA deficiency also has been previously shown to increase circulating lactate concentrations consistent with increased glycolytic flux. In prior studies, whole-body knockout mice exhibited exercise-induced hyperlactatemia [3]. In addition, MondoA has been shown to be lactate and pH-responsive, suggesting lactatemia could be part of a negative feedback loop controlling glycolytic flux [4, 5]. However, the phenotype of generalized MondoA knockout (gKO)^1^ mice in the context of chronic caloric excess has not been determined.

Futile cycling is a rather poorly understood process that likely participates in dynamic control of whole-body energy homeostasis, by tightly regulating opposing metabolic pathways [6]. Perhaps best known in the context of adaptive thermogenesis, energetic uncoupling of the electron transport chain in brown adipocytes via uncoupling protein 1 (UCP1) regulates body temperature and plays a role in whole body energy balance. Recently, additional forms of futile cycling for adaptive thermogenesis, e.g. futile creatine cycling, have been defined in beige and white adipocytes [7, 8]. Beyond heat production, less is known about “multi-organ” or systemic futile lipid cycles, such as triglyceride-fatty acid [6, 9, 10] and glycerol-glucose cycling [11], where ATP is consumed in the process of lipolysis/re-esterification or sequestering excess glucose. It is tempting to speculate that such futile cycles respond to environmental and nutritional cues to maintain systemic energy and metabolic balance.

To understand the potential role of the nutrient-regulated transcription factor MondoA in the control of whole-body metabolism, and as a step towards assessing this factor and its downstream targets as candidate therapeutic targets, we developed MondoA gKO mice. Given the previously described role in cellular glucose homeostasis and lipid storage, gKO mice were carefully phenotyped for measures of metabolic health at baseline and in the context of chronic caloric excess. The gKO mice were resistant to diet-induced obesity with enhanced glucose tolerance and insulin sensitivity and reduced hepatic steatosis. gKO mice exhibit increased whole body energy expenditure in the absence of adipose browning. Signatures of enhanced glucose uptake and flux together with evidence of adipocyte triglyceride lipolysis/re-esterification strongly suggest that futile cycles are driving the observed energy expenditure. These results add to a growing recognition of energy-expending futile cycles in the control of whole-body energy homeostasis and identify potential new therapeutic avenues for diseases of altered nutrient balance.

## Material and methods

### Animal studies

A generalized *Mlxip* (MondoA) deletion mouse line was created by breeding the MondoA-floxed line [1] to mice ubiquitously expressing EIIa-Cre recombinase (Jackson Laboratory #003724) maintained on a C57Bl/6NJ (Jackson Laboratory #005304) background. The adipose-specific KO was created using Adiponectin-Cre (Jackson Laboratory #028020) [12]. Mice were housed in a facility under a 12-hour light/12-hour dark cycles. Mice were maintained on standard chow (Lab Diet 5010) or a 60% fat HFD (Research Diets, D12492i) for 16 weeks. For tissue collection, mice were fasted for 4 hours prior to anesthesia with intraperitoneal pentobarbital (Sagent, 100 mg/kg). Experiments were conducted on groups of 6– 10 mice with 2-3 independent experiments.

### Metabolic profiling

Glucose and insulin tolerance testing (GTT and ITT, respectively) was conducted after a 5-hour fast, as previously described.[1] Tail tip blood glucose measurements were obtained using a handheld glucometer (AlphaTrak2 #71681-01) immediately prior to and 15, 30, 45, 60, and 90 minutes following intraperitoneal glucose (1 g/kg) or insulin (0.75U/kg) injections. Pyruvate tolerance testing (PTT) utilized the identical timepoints, following 1 g/kg intraperitoneal injection of sodium pyruvate (pH 7.3-7.5, 0.22μM filter sterilized). Plasma insulin was measured using Mouse Insulin ELISA kit (Mercodia). Cholesterol and triglycerides were measured using Infinity colorimetric assays (ThermoScientific), and blood lactate was obtained using a blood lactate measuring meter (NovaBiomedical). *In vivo* stimulated lipolysis assay was performed in fed mice. Tail blood sample was collected via collection tubes at baseline, then at 10 and 30 minutes after administration of CL316,243 (Sigma C5976) at 0.1ng/kg via intraperitoneal injection. Non-esterified free fatty acid (FFA) and glycerol in serum were measured using HR Series NEFA-HR colorimetric assays (Wako, Fujifilm) and Glycerol colorimetric assay kit (Cayman Chemical), respectively. For body composition measurements, mice were gently restrained in a tube while body composition was measured in an EchoMRI 100 analyzer (EchoMRI, Houston TX). Data was visualized using CalR2 [13].

### Indirect calorimetry

Metabolic cage experiments were performed with a Promethion CORE metabolic cage system (Sable Systems International, North Las Vegas, NV). Mice acclimated in the Promethion metabolic cages for 72 hours at 22°C before a 24-hour recording period. Oxygen consumption (VO_2_) and carbon dioxide production (VCO_2_) were measured by indirect calorimetry every 5 minutes and used to calculate the respiratory exchange ratio (RER = VCO_2_/VO_2_) and energy expenditure using the Weir equation (Energy expenditure = 3.941 kcal/L x VO_2_ + 1.106 kcal/L x VCO_2_) [14]. Food intake, water intake, and body mass were measured in the metabolic cages gravimetrically, and ambulatory activity was recorded through a beam break array.

### Cold tolerance testing

Male mice age 10-12 weeks were housed in thermoneutral incubators (28-30°C) for 14 days. Mice were transferred to pre-chilled at 4°C, single housing, without bedding, food, or water. Rectal core temperature was measured prior to cold exposure, and every 60 minutes while >35°C, every 30min <35°C, and every 15 min <30°C. Animals were removed from the study at core temperature <25°C.

### Histology

Paraffin-embedded adipose tissue was prepared as previously described including overnight fixation in 4% paraformaldehyde in PBS and dehydration through sequential ethanol washes [15]. The tissues were then embedded in paraffin, sectioned at 10μm, and stained with hematoxylin and eosin (H&E) or perilipin (CS 3470, 1:500) by the IDOM Histology Core. For adipose tissue, following deparaffinization, slides were subjected to heat antigen retrieval in a pressure cooker with Bulls Eye Decloaking buffer (Biocare). Frozen liver tissue was embedded in Tissue-Plus OCT (Fisher) and sectioned by the University of Pennsylvania Molecular Pathology and Imaging Core, prior to Oil Red O staining. Slides were imaged by the Children’s Hospital of Philadelphia Pathology Core using Aperio ImageScope (Leica Biosystems).

### RNA isolation and qRT-PCR

Total RNA was isolated from frozen tissues or cells using the RNeasy Mini Kit (QIAGEN) and treated with RNase-Free DNase Set (QIAGEN) per manufacturer’s instructions. cDNA was synthesized using the Affinity Script cDNA Synthesis Kit (Agilent Technologies) or High-Capacity cDNA Reverse Transcription Kit (ThermoFisher Scientific #4368814) with 0.25 μg total RNA. PCR reactions were performed using Brilliant III Ultra-Fast SYBR Green QPCR Master Mix (Agilent Technologies) on a QuantStudio 6 Flex Real-Time PCR System (Applied Biosystems) or TaqMan Fast Advanced Master Mix (Applied Biosystems) on a CFX384 Real Time System (BIO-RAD) system with specific primers for each gene (Table S1). Target mRNAs expression was normalized to *Rplp0* (36B4) or *Tbp*. All experiments were repeated independently, with samples in either triplicate or quadruplicate.

### RNA-Seq library preparation, sequencing and analysis

GENEWIZ, LLC. (South Plainfield, NJ) performed RNA library preparations and sequencing reactions per standard operating procedure [16, 17]. Reference transcripts, e.g. GRCm38 (mm10), were counted with Salmon v1.10.2 [18]. The counts were summarized at gene level and differentially expressed genes were identified with DESeq2 1.40.2 [19]. Differentially expressed genes were defined by a false discovery rate below 0.05 and 1.2-fold expression change (up or down). gProfiler web service was used for pathway enrichment analysis, using with FDR < 0.05 cutoff and enrichment > 2 [20].

### Mouse adipose-derived cell isolation

Adipocytes and stromal vascular cells were isolated as previously described [21].

### Human adipose-derived stem cell (hASC) culture and differentiation

hASCs (PT-5006, batch #21TL104467, Female, Black, age 55 from Lonza Walkerville Inc, Walkersville, MD; or ASC-F, lot# ASC080816A, Female, Black, age 58 from Zen-Bio, Durham, NC) were incubated at 37°C / 5% CO_2_ for 5-6 days in ‘Media 22’ (225 ml DMEM (Gibco #11885-054), 225 ml F-12 (Gibco #11765-054), 50 ml Fetal Bovine Serum, heat-inactivated (Sigma #F4135), 5 ml Penicillin-Streptomycin 10,000 U/ml (Gibco #15140-122), 2.5 ml Amphotericin B (Gibco #15290-018), 0.5 ml Gentamicin 50 mg/ml (Gibco #15750-060)). To differentiate, 0.8e6 per T150 collagen flask (Corning Biocoat collagen #354486) were incubated for 3 days, followed treatment with Preadipocyte Growth Medium (PromoCell GmbH, Heidelberg, Germany, #C-27410) with Supplement Mix (C-39425). On day 4, media was replaced with Adipocyte Differentiation Medium (DM2) (ZenBio Durham, NC #DM2-500), termed day 0 of differentiation. DM2 media was refreshed with 2/3 volume on day 4. On day 7, media was replaced with DMAM (50/50 mixture of DM2 and Adipocyte Maintenance Medium (AM1), ZenBio Durham, NC #AM1), and changed every 3-4 days following. Post differentiation assays conducted on day 16-17.

### CRISPR Knock-Down in hASCs

Day 9 differentiated hASCs used for Alt-R Cas9 enzyme mediated knock-down (KD) of target genes with SE Cell Line 96-well Nucleofector X Kit (Lonza V4SC-1960), Alt-R S.p. Cas9 Nuclease V3 62 μM (Integrated DNA Technologies, (IDT), #1081059), Alt-R Cas9 Electroporation Enhancer 100 μM (IDT, #1075916), sgRNAs 100 μM (IDT custom-made, see Table S2). Trypsinized and PBS-washed cells were resuspended in SE Cell Line Nucleofector Solution with Supplement 1 reagent at ∼7.2-7.4e6 cells/ml based on pre-spin cell count. Alt-R S.p. Cas9 Nuclease V3 and sgRNA guide combined at 1:1.45 ratio in PBS (total 10 μl) for 15min at room temperature was added to 2 μl Alt-R Cas9 Electroporation Enhancer 100 μM, IDT #10007805) and 53 μl of cells. 25 μl of cell/ Cas9/sgRNA mixture was plated in 96 well nucleocuvette plate, followed by nucleofection on 4D Nucleofector and 96-well shuttle units (Lonza) with program code 96-DS-150 cell line SE. 75 μl DMAM medium was added, and 85 μl (∼125,000 cells) was transferred to 96 well collagen coated plates (Corning BioCoat #354407) preloaded with 65 μl DMAM medium. Cells were grown until differentiation day 16-17. For CRISPR hASC Seahorse assays, cells were plated at 20,000-30,000 cells per 96-well and for Fatty Acid Oxidation assays at 125,000 cells per 96-well. To confirm INDEL knockout, isolated genomic DNA from cells 4-5 days post CRISPR was sequenced using custom genomic PCR and sequencing primers (Table S3) from IDT with the Expand High Fidelity PCR System (Millipore #11732650001). INDEL KO efficiency was determined using Synthego’s ICE CRISPR Analysis Tool (<u>synthego.com/products/bioinformatics/crispr-analysis</u>) with the non-CRISPR genomic DNA sequence as reference control. Knockdown was further confirmed by western blot (for MondoA and Txnip) or QT-PCR (for Arrdc4, given no satisfactory antibody available).

### Western blotting

Transfected cells were incubated on ice for 20 minutes with TNET lysis buffer (50mM Tris-HCl (pH7.4), 150mM NaCl, 2mM EDTA, 1% Triton X-100, 0.5% cholate) and diluted 100X Halt Protease/Phosphatase inhibitor cocktail, EDTA free (Thermo #78445). Lysate was sonicated at 20 Hz for 3 seconds with microtip followed by centrifugation at 9,600 x g for 10 minutes. Protein quantification of supernatant was performed with BCA protein assay kit (ThermoFisher Scientific #23227). Gel electrophoresis was performed with 4-12% NuPage Bis-Tris gels (Invitrogen WG1402) and MOPS running buffer (Invitrogen NP0001, and gel electroblotted onto nitrocellulose membrane (Invitrogen LC2009) using Transfer buffer (Invitrogen NP00061) and 10% methanol. Blots were blocked using Odyssey Intercept TBS blocking solution (LiCor 927-60001) and probed with 1:1000 MondoA, Txnip, ATGL, and 1:2000 B-actin antibodies (Table S4) in Intercept antibody diluent-Tween/TBS (LiCor 927-65001), followed by secondary detection antibody (1:10,000 dilution) with donkey anti-rabbit IRDye 800 (Odyssey 626-32213) and donkey anti-mouse IRDye 680 (926-68022). Washes were conducted with TBS + 0.05% Tween20. Blots were scanned on Odyssey CLx and detected proteins were quantified using Image Studio software 5.2 and normalized to relative B-actin levels and HPRT1 control.

### *In vitro* isoproterenol-stimulated lipolysis assay

For stimulated lipolysis assay, Day 7 differentiated hASCs were replated at 65,000 cells/well in collagen 96 well plates in DMAM media. Cells were differentiated to Day 14. On the day before assay, media replaced with “starvation media” (DMEM/F12 (Gibco #11330-032) plus 0.1% BSA (Sigma BSA A7888)). Cells were washed with PBS (without magnesium or calcium) and treated with 10 nM isoproterenol-HCl (ISO) (Sigma I6504) in 100 μl “Assay buffer” [117 mM NaCl, 4.7 mM KCl, 2.5 mM CaCl, 1.2 mM MgSO_4_, 24.6 mM NaHCO_3_, 5 mM HEPES, pH 7.4 supplemented with 10 mM glucose, 1% fatty acid free, BSA (Roche #3117057011)] for 2h at 37°C / 5% CO_2_. As indicated, cells were pre-treated with DMSO 0.1% control or Triascin C (Sigma T4540) and Etomoxir (EMD Millipore #236020) resuspended in DMSO at 10 mM and diluted with LAB for final assay of 10 μM for 1h at 37°C / 5% CO_2_, then dose with 10 nM ISO in final 100 μl volume for 2h at 37°C / 5% CO_2_. Supernatant was collected for FFA [HR Series NEFA-HR(2), FujiFilm Wako Pure Chemical Corp] and glycerol (Free Glycerol Detection Reagent, F-6428 Sigma) release measurements. Cells were washed in PBS and stained with Hoechst 33342 trihydrochloride (Invitrogen H3570) at 1:2000 in PBS for 30’ at room temperature. Nuclei were counted on a BioTek Cytation 5 (Agilent) for data normalization.

### Seahorse XF Cell Stress Test

CRISPR hASC KD cultures (20,000 cells/well) plated onto rat collagen type I coated plates (Advanced BioMatrix #5056) were assessed for effect on glycolytic function/extracellular acidification rate (ECAR) assays using Seahorse XFe96 Pro Glycolysis Stress Kit (Agilent #103020-100) per manufacturer protocol with Seahorse XFe96/XF Pro cell culture microplates (Agilent #103794-100), XF Calibrant (Agilent #100840-000), Seahorse XFe96 Extracellular Flux Assay kits (Agilent), assay medium composed of XF DMEM medium, pH 7.4 (Agilent #103575-100), Seahorse XF L-Glutamine 2 mM (Agilent #103579-100). Final concentrations of glucose 10 mM, oligomycin 1 μM, and 2-DG 50 mM were used in the assay. To assess for effect on mitochondrial respiration/oxygen consumption rates (OCR), CRISPR hASC KD cultures (30,000 cells/well) plated onto rat collagen type I coated plates (Advanced BioMatrix #5056) underwent mitochondrial respiration assays using Seahorse XFe96 Pro Mitochondrial Stress Kit (Agilent #103015-100) per manufacturer protocol with Seahorse XFe96/XF Pro cell culture microplates (Agilent #103794-100), XF Calibrant (Agilent #100840-000), L-carnitine (Agilent #103689-100), Seahorse XFe96 Extracellular Flux Assay kits (Agilent), assay medium composed of Seahorse XF DMEM medium, pH 7.4 (Agilent #103575-100), Seahorse XF Glucose 10 mM (Agilent #103577-100), Seahorse XF Pyruvate 1 mM (Agilent #103578-100), Seahorse XF L-Glutamine 2 mM (Agilent #103579-100). Final concentrations of oligomycin 1.5 μM, FCCP 1 μM, and Rotenone + Antimycin A 0.5 μM used in the assay. Data was normalized to nuclei counts, as above, using WAVE analysis software to determine ECAR (glycolysis, glycolysis capacity, glycolysis reserve) and OCR (spare respiratory capacity and maximum respiration).

### Fatty Acid Oxidation Assay

Palmitic acid (PA) was conjugated with BSA by adding ^3^H-PA ([9,10-^3^H(N)]-PA, 1mCi (37MBq), Perkin Elmer NET43001MC) to 3 ml of pre-warmed PBS in glass vials at 60°C to generate PA/PBS mixture, then added to 3 ml of fatty acid free BSA (4 mg/ml) in PBS preheated to 42°C to generate 4X ^3^H-PA 20 μCi/ml stock solution. Prior to the assay, hASC were treated with “starvation medium” for overnight incubation at 37°C / 5% CO_2_. For the assay, one part of 4X ^3^H-PA/FAF-BSA was mixed with 3 parts 1.33X substrate mixture [1 M L-carnitine (6 ul), 1 M glucose (15 μl), 1 M HEPES (0.12 ml), 5X KHB (1.2 ml: 555 mM NaCl, 23.5 mM KCl, 10 mM MgSO4 in H2O, pH 7.4 with 6N HCl), ddH_2_O (3.17 ml), 6N NaOH (3.75 μl)] at 37°C. Cells were washed with PBS, and treated with 10 or 20 μM Etomoxir, 1 μM FCCP, 0.1% DMSO in above “assay buffer” for 1h at 37°C / 5% CO_2_. PBS-washed twice, cells were exposed to 1X ^3^H -PA/FAF-BSA mixture with 10 or 20 μM Etomoxir, 1 μM FCCP, 0.1% DMSO for 4h at 37°C / 5% CO_2_. Supernatant was collected for TCA precipitation by adding equal amount of 10% cold TCA, shaking at room temperature for 10’, centrifuging 870 x g for 10’, and combining supernatant mixture (90 μl) with 6N NaOH (14.3 μl). Mixture was transferred to prepared filter plate (0.2 ml ion exchange resin slurry per well, spin at 870 x g for 5 minutes), and centrifuged at 870 x g for 5 minutes at room temperature. Flow -through was mixed with equal volume OptiPhase HiSafe 3, and ^3^H cpm counts were obtained on Perkin Elmer Microbeta2. Data normalized to HPRT1 control or DMSO control.

### 2-Deoxyglucose uptake

Adipocytes plated in 96 well plates were incubated overnight in DMEM/F12 with 0.5% BSA (Sigma A7888) and penicillin/streptomycin. The following morning, cells were washed and incubated with assay buffer [Kreb’s Ringer Phosphate buffer with 0.1% BSA and 0.22 mg/ml sodium pyruvate (Sigma P-2256)] for 2 hours at 37°C. Cells were treated with 1nm Insulin (Sigma I9278) for 30 minutes at 37°C, followed by 1-^14^C 2-Deoxyglucose (Revity, NEC495A) diluted 1:10 in 1.1 mM cold 2-Deoxyglucose (Sigma D6134) for 1 hour at 37°C. At the end of the assay, media was removed and cells were washed with assay buffer. Cells were lysed with 1% Triton X-100 in PBS and added to Deepwell Luma Plates, dried down overnight on a heat block at 42°C, and read in a Wallac MicroBeta Counter.

### Glucose consumption and lactate production

The day following Crispr-Cas9 mediated deletion of HPRT, MondoA, Txnip or Arrdc4, media was changed to fresh DMAM media. Media glucose and lactate concentrations were measured using an ADVIA Chemistry XPT System (Siemens).

### Palmitate and acetate uptake assays

For palmitate assays, adipocytes plated in 24 well plates were incubated overnight in DMEM/F12 media with 0.1% BSA (Sigma A7888). The next morning, adipocytes were treated for 1hr at 37°C with 10nm isoproterenol diluted in media or LAB assay buffer. 0.25 μCi of 1,2-^14^C–acetic acid (American Radiolabeled Chemicals 0173A) or U-^14^C-palmitate (Perkin Elmer NEC534250) were added to each well and incubated at 37°C for 2 hours. Plates were then placed on ice, washed with ice-cold PBS, and frozen at –20°C overnight to assist with lysis. The following morning, 125 μL of Mammalian Protein Extraction Reagent (Thermo 78505) was added to each well and plates were shaken for 1 hour at room temperature to induce lysis. Lysates were transferred to individual 1.5 mL tubes, and each well was rinsed with 175 μL of PBS and subsequently added to each tube. 450 μL of chloroform:methanol (1:1, v/v) was added, samples were vortexed vigorously for 30 seconds and centrifuged at 20,000 × g for 5 minutes at room temperature. The bottom organic layer (25 μL) was transferred to a 7 mL glass scintillation vial and 6 mL of Optiphase Hisafe (Perkin Elmer) was added. Vials were vortexed briefly and each vial was added to a scintillation counter for screening for ^14^C emission.

### Statistics

A Student’s *t* test was performed for normally distributed 2-group comparisons, based on a Shapiro-Wilk test (α = 0.05). Multiple comparisons were analyzed by 1-way ANOVA with Tukey’s post hoc test, or 2-way ANOVA with Tukey’s post hoc test, as indicated. Graphing and statistical analysis utilized GraphPad Prism 8.03 (GraphPad Software).

### Study approval

All animal experiments were approved by the Institutional Animal Care and Use Committees of the Perelman School of Medicine at the University of Pennsylvania and of Pfizer Inc. and conducted in accordance with local, state, federal guidelines and the Guide for the Care and Use of Laboratory Animals from the National Institutes of Health.

## Results

### MondoA gKO mice are resistant to diet-induced obesity with decreased adiposity

The MondoA gKO mouse line was derived by crossing mice harboring the previously published floxed allele[1] to mice with germline Cre expression (EIIa Cre; Figure S1A) and knockout was confirmed by QT-PCR in multiple tissues (Figure S1B). Expression of the two key MondoA target genes *Txnip* and *Arrdc4* was substantially depressed in gKO mice, particularly in skeletal muscle and heart where its expression is enriched, but not in liver, where MondoA is not highly expressed. Mice were born in expected Mendelian ratios and developed normally as has been described [3], though male gKO mice were sterile (data not shown). To assess the effect of global MondoA deficiency in the context of caloric excess, cohorts of gKO and wildtype (WT) littermates were challenged with 60% high fat diet (HFD) compared to chow diet (CD) for 16 weeks. gKO mice were protected from diet-induced obesity (Figure 1A), which was not observed in the muscle-specific KO (msKO, [1]). When assessed by echoMRI, this roughly 25% reduction in body weight was attributable to decreased fat mass (Figure 1B). Lean (muscle) mass in gKO HFD mice was similar to both WT and gKO CD cohorts, and significantly less than WT HFD littermates. To confirm these whole-body measurements, adipose depots were surgically excised and weighed. Weight of the brown adipose (BAT) and inguinal (or subcutaneous) white adipose (iWAT) depots were reduced by nearly half in the HFD gKO mice compared to HFD WT, while epididymal (or visceral) adipose (eWAT) mass was unchanged (Figure 1C).

**Fig 1.**
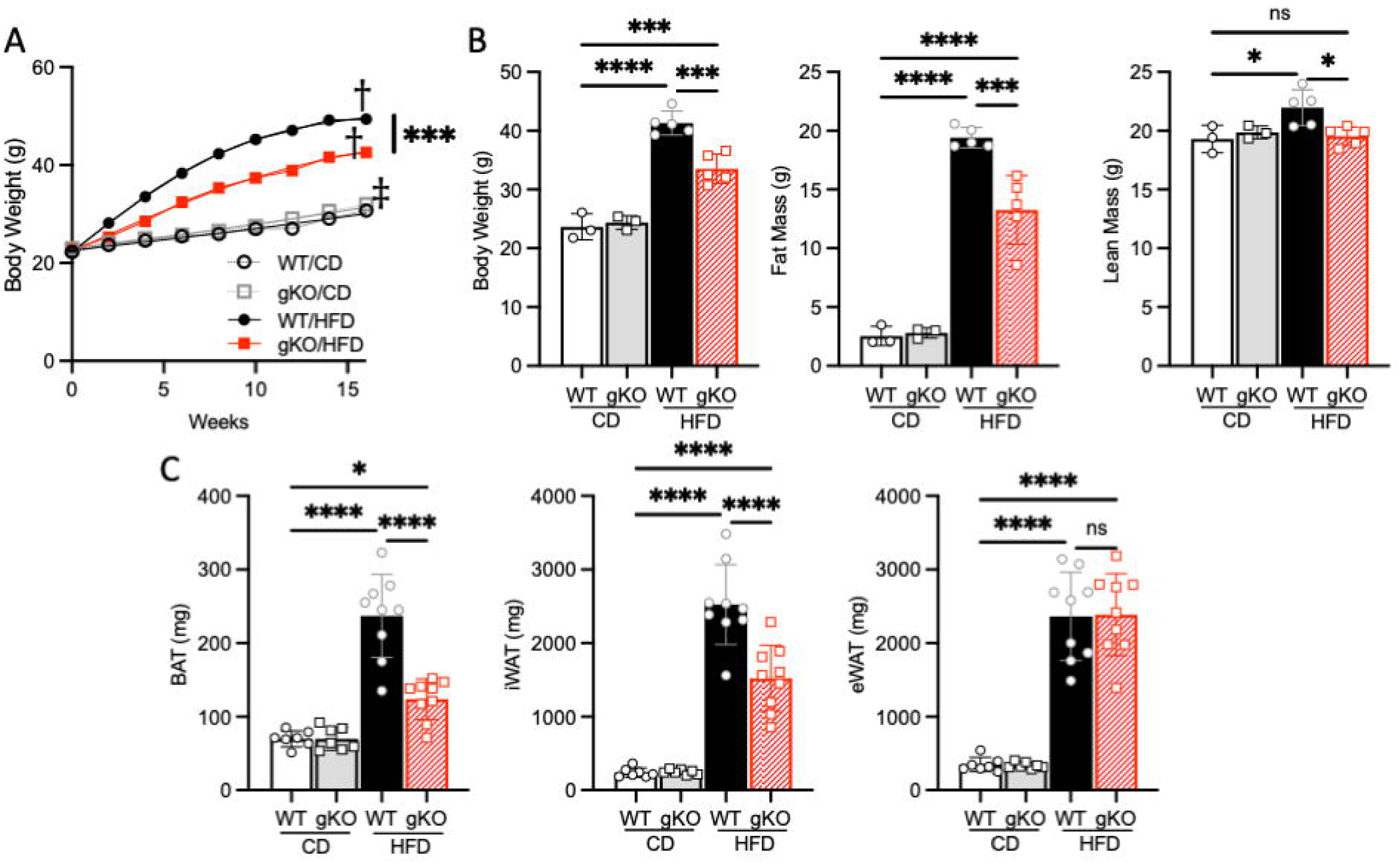
Whole-body MondoA deficiency protects against diet-induced obesity with reduced adiposity. A) Body weight trends for WT and gKO MondoA mice on chow and 60% kcal HFD over sixteen weeks (n=14-25). B) MRI body composition data (n=3-5) and c) gross weights of brown (BAT), inguinal white (iWAT) and epididymal white (eWAT) adipose tissues (n=7-9). CD, chow diet; HFD, high fat diet; KO, knockout; WT, wildtype. P values * < 0.05, *** <0.001, ****<0.0001, ns, not significant displayed on graphs; ‡ < 0.05 and † < 0.001 compared to WT CD. The data represent mean ± SD. All statistical significance determined by 2-way ANOVA with Tukey’s multiple-comparisons post hoc test.

We next assessed glucose handling and lipid storage in the context of the HFD. The effect of MondoA deficiency in the setting of HFD resulted in improved fasting blood glucose levels, insulin sensitivity and glucose tolerance (Figure 2A-C), comparable to the msKO model [1]. As expected, the leaner gKO mice had a normalization of hyperinsulinemia (Figure 2D, compared to WT HFD littermates) and improved measures of circulating cholesterol and triglycerides. No differences were observed comparing the CD WT and gKO littermates. The leaner phenotype of the gKO mice also corresponded with reduced neutral lipid accumulation in the gKO HFD liver (Figure S2A). As predicted by the gross weight changes, BAT and iWAT from gKO HFD mice exhibited reduced lipid droplet size compared to the WT control group (Figure S2B). Perilipin staining was similarly reduced in the gKO HFD BAT. Lipid droplet size and inflammatory infiltration in eWAT appeared similar between both genotypes. Notably, the MondoA target *Arrdc4*, but not *Txnip*, was highly regulated in all adipose depots (Figure S3).

**Fig 2.**
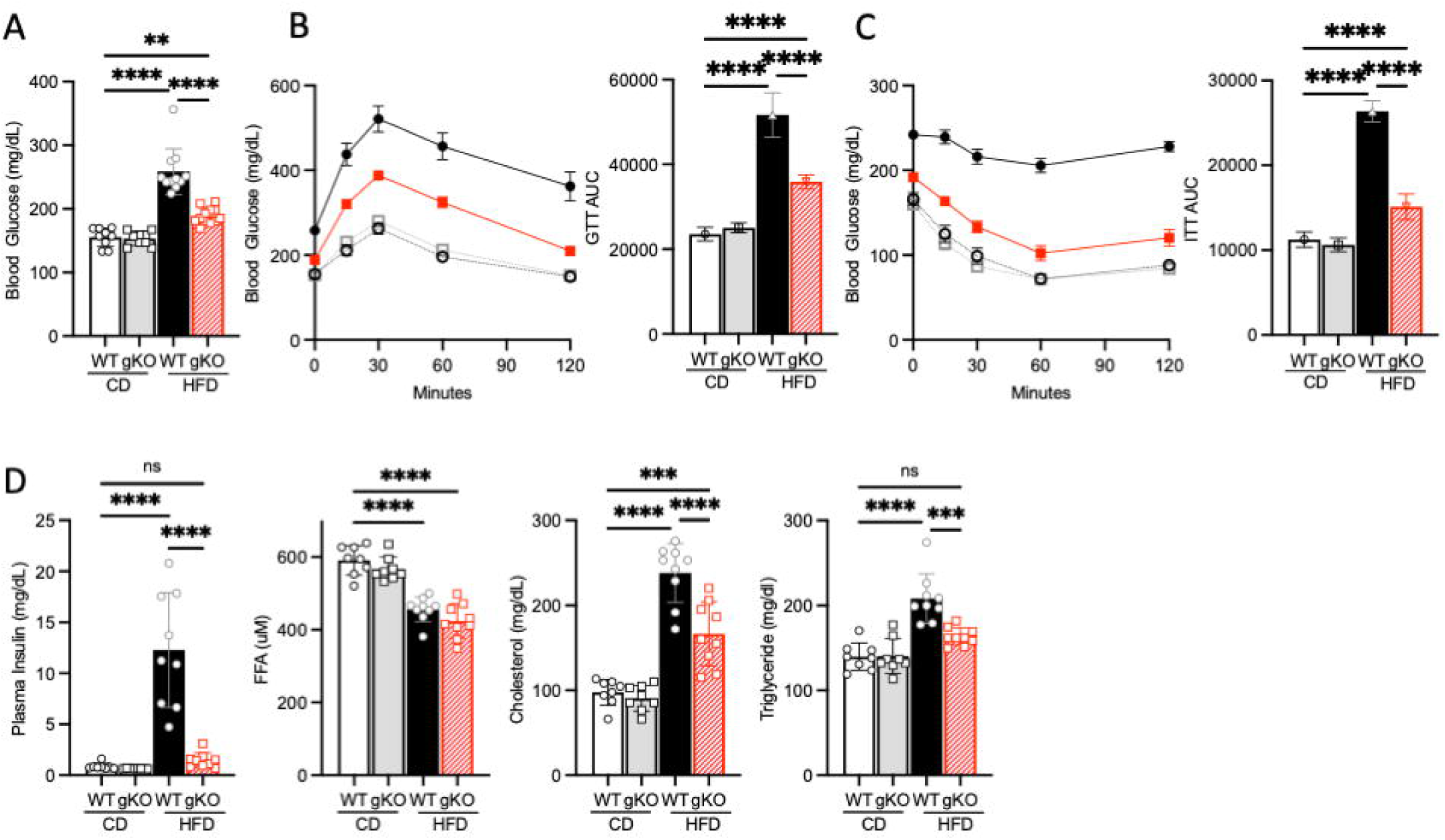
MondoA gKO mice have improved glucose tolerance, insulin sensitivity, and reduced dyslipidemia. A) Fasting whole blood glucose in WT and gKO male littermates on chow and HFD for 10 weeks (n=8-12). B) Glucose and C) insulin tolerance tests (left panel) and area under the curve (right panel). D) In the same cohort of mice, fasted plasma measurements of insulin, free fatty acids (FFA), total cholesterol, and triglycerides. CD, chow diet; HFD, high fat diet; KO, knockout; WT, wildtype. Data displayed as mean ± SEM. Two-way analysis of variance (ANOVA) followed by multiple-comparisons test. P values * < 0.05, **< 0.01, *** <0.001, ****<0.0001 displayed on graphs.

### MondoA global deficiency results in increased energy expenditure and protection against cold exposure without evidence of BAT or WAT respiratory uncoupling

To further parse the basis for the MondoA gKO protection from diet-induced obesity, littermate WT and gKO mice were subjected to indirect calorimetry. HFD-fed cohorts had increased oxygen consumption and decreased dark cycle RER (Figure 3A-B). Adjusting for lean mass, energy expenditure was increased in the gKO HFD cohort (Figure 3C). The leaner gKO HFD mice were comparatively more active than the WT HFD cohort and consumed the same cumulative calories as the chow fed cohorts (and more than the WT HFD group) despite the lean phenotype (Figure 3D-E). We next sought to assess the thermogenic function of the gKO mice. For these studies, chow-fed WT and gKO littermates were challenged with cold exposure following 14 days of thermoneutrality. MondoA gKO mice had significantly greater ability to defend their core body temperature over five hours compared to control mice (Figure 4A). Surprisingly, however, gene expression markers of classical non-shivering thermogenesis were not activated in the gKO BAT (Figure 4B).

**Fig 3.**
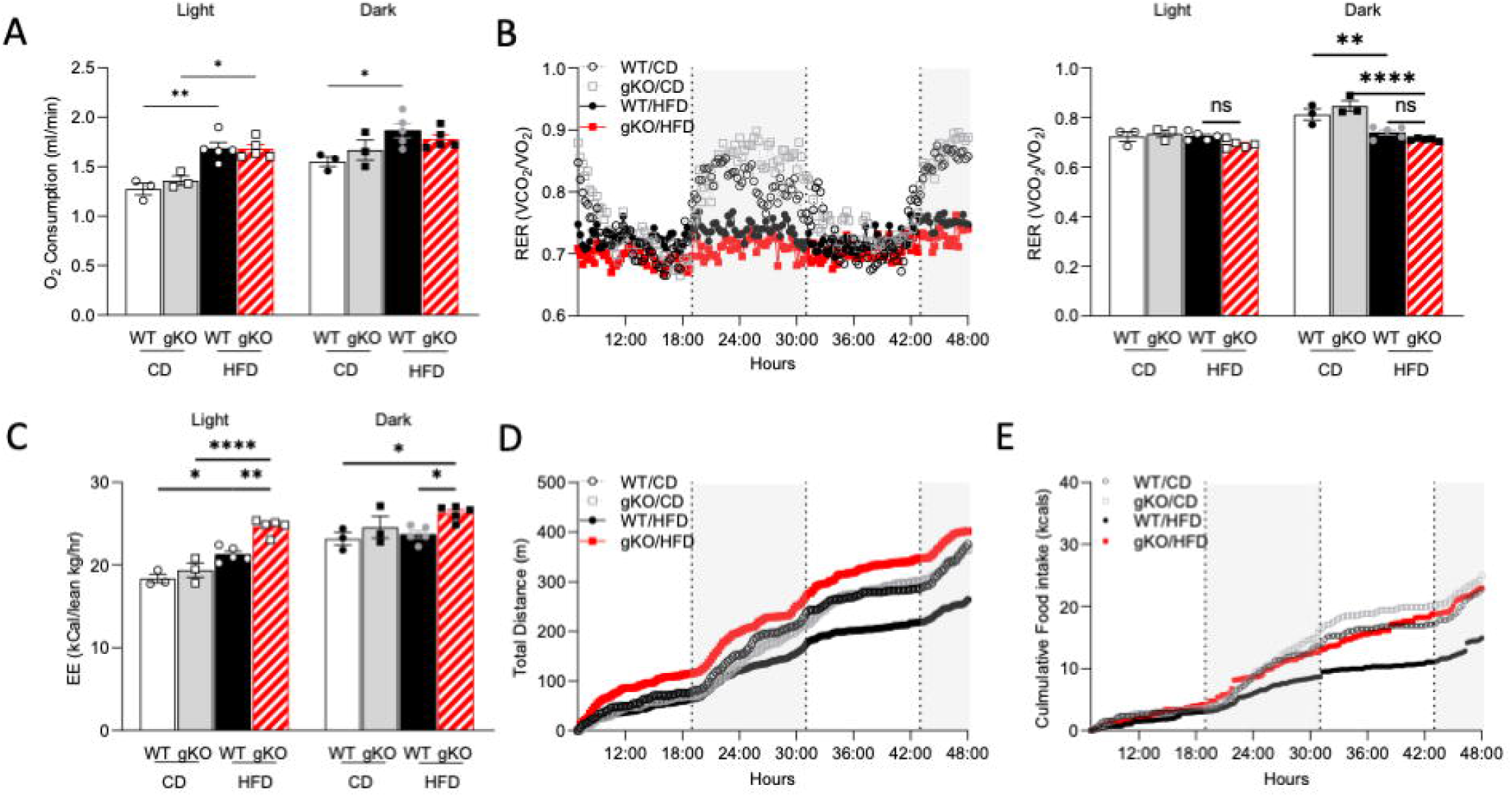
Lean HFD-fed gKO mice have increased energy expenditure despite higher caloric intake. Chow and HFD-fed WT and gKO male littermates were analyzed in metabolic cages (n=3-5). A) Oxygen consumption in light and dark cycles (averaged over 48 hours). B) Respiratory exchange ratio (RER) over time (left panel) and average by light and dark cycles (right panel). C) Total energy expenditure normalized to lean body mass as measured by MRI in Fig. 1B. D) Cumulative distance traveled and E) food consumed over 48 hours. CD, chow diet; HFD, high fat diet; KO, knockout; WT, wildtype. Data displayed as mean ± SEM. Two-way analysis of variance (ANOVA) followed by multiple-comparisons test (within specific light cycle). P values * < 0.05, **< 0.01, *** <0.001, ****<0.0001 and ns, not significant displayed on graphs.

**Fig 4.**
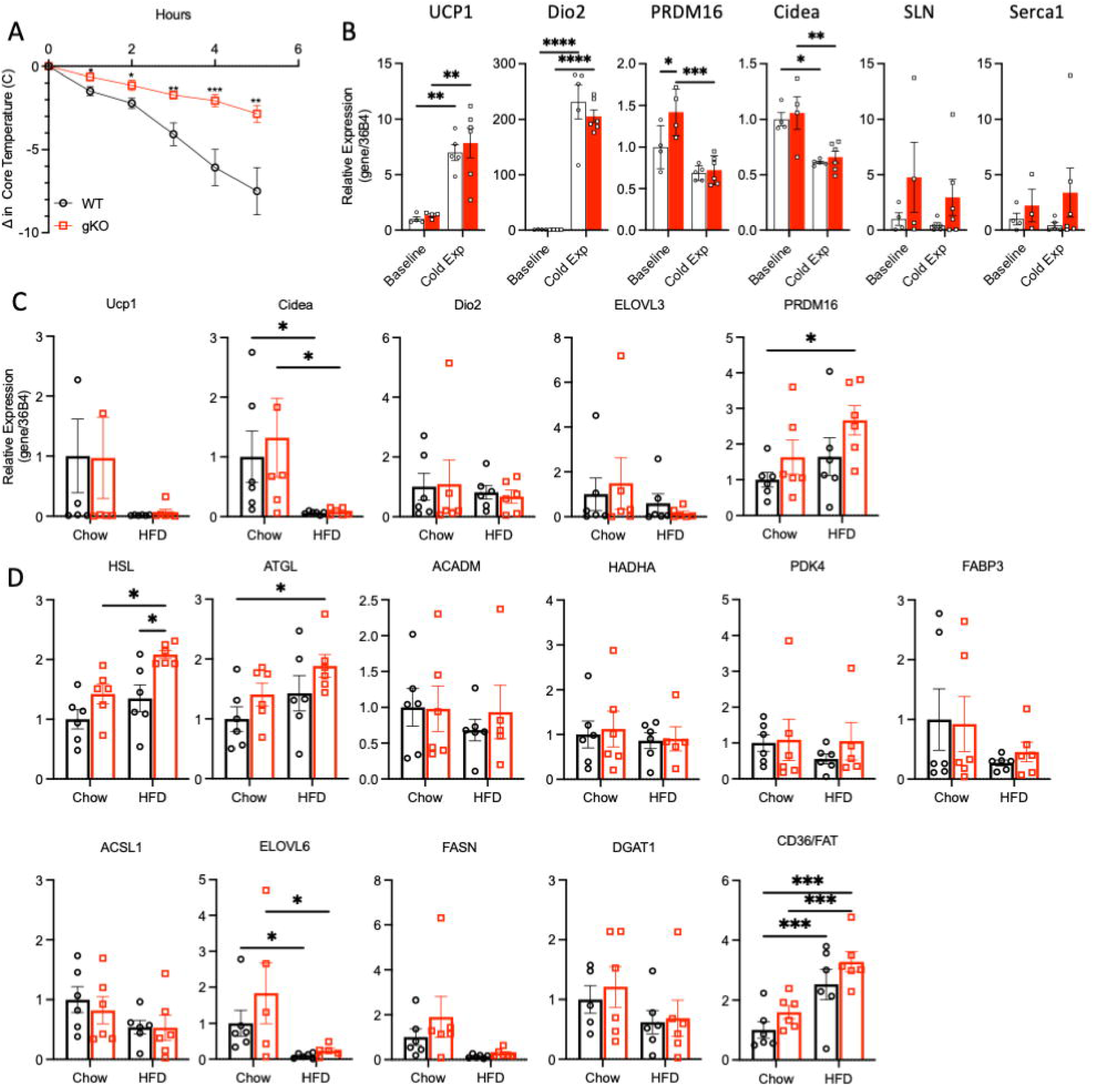
MondoA gKO mice have increased cold tolerance without evidence of increased UCP-1 expression in BAT and lipolytic signature without beiging in iWAT. A) Chow-fed WT (n = 16) and MondoA gKO (n = 18) mice age 10-12 weeks were subjected to thermoneutrality for 14 days followed by acute cold exposure (4 °C). Core rectal temperature was monitored over a 5-h period. The change in core temperature ± SEM is shown in the graph as a function of time. B) Quantitative real-time RT-PCR performed from RNA isolated from BAT for targets of classical and nonclassical non-shivering thermogenesis. The values represent mean arbitrary units normalized to a 36B4 transcript (control). QT-PCR targets for C) non-shivering thermogenesis and D) lipid synthesis, storage and utilization from iWAT isolated from chow and HFD-fed WT and MondoA gKO male littermates (n=5-6). Data displayed as mean ± SEM. Two-way analysis of variance (ANOVA) followed by multiple-comparisons test. P values * < 0.05, **< 0.01, *** <0.001, ****<0.0001 displayed on graphs.

To further explore the gKO thermogenic response, we performed bulk RNA sequencing (RNAseq) on BAT from normothermic chow and HFD-fed animals. Principal component analysis (PCA) demonstrated that the cohorts segregated mainly by diet (Figure S4A). Analysis of differentially regulated gene expression between CD-fed littermates solely identified the downregulation of *Arrdc4*. In HFD-fed mice, the expression of one gene was upregulated and 16 genes were downregulated in gKO mice compared to WT littermates. Kyoto Encyclopedia of Genes and Genomes (KEGG) pathway analysis was notable for downregulation of multiple inflammatory pathways (Figure S4C), and no upregulated pathways. Target gene *Arrdc4* was significantly downregulated in BAT. Genes involved in immune system activation were also significantly repressed by MondoA deletion, including Tnf, with downward trends in several other markers (Figure S4D). The downregulation of the proinflammatory signature was likely caused by the lean phenotype of MondoA deficiency, given that inflammatory cells including macrophages have been known to infiltrate to obese adipose tissue [22, 23]. Taken together, despite the greater tolerance to cold exposure, the BAT tissue of the gKO does not display significant gene expression changes including a lack of induction of genes involved in respiratory uncoupling.

Given the absence of gene expression signatures in BAT to explain increased energy expenditure, despite impressive activation of adaptive thermogenesis, we explored other tissues capable of contributing to the thermogenic response. In isolated gastrocnemius, there was a modest (30%) increase in *Serca1* expression but not in other markers of non-shivering thermogenesis (Figure S5A). Furthermore, the msKO HFD model had not previously shown robust protection from obesity [1], suggesting an alternative tissue(s) contributing to the thermogenic response. Given that the capacity for white adipocyte “browning” is a well-described response to stimuli as varied as cold exposure, exercise, and aging, we next measured expression of beiging-related and metabolic genes in eWAT. *Prdm16* appeared elevated in HFD gKO mice, but other known markers were not significantly modified (Figure 4C) [24]. Hormone-sensitive lipase (*Hsl*) and adipose triglyceride lipase (*Atgl*) were notably elevated, suggesting an increased lipolytic state. Other genes involved in lipid synthesis, storage and oxidation that might explain the lipodystrophic changes were not differentially expressed, with the exception of *Cd36* (Figure 4D). To further pursue these observations in an unbiased way, bulk RNAseq was performed on iWAT tissue. MondoA (*Mlxip*) and *Arrdc4* were downregulated, as anticipated. PCA demonstrated the predominant effect of diet relative to genotype (Figure S4B, top). Factor-based analysis of DEGs indicated 6 upregulated and 5 downregulated genes due to KO (Figure S4B, bottom). No downregulated pathways were identified by KEGG analysis; upregulated pathways were exclusively focused on mineral absorption/inorganic ion regulation (driven by *Slc4a1, Hcrtr1, Mt2, Mt1*).

### Absence of MondoA glucose uptake “brake” promotes enhanced glycolytic flux

Previous work has shown that MondoA deficient mice develop lactatemia, consistent with a role for the loss of appropriate inhibition of glucose uptake by MondoA-regulated Txnip [3–5]. We confirmed that the gKO mice exhibited lactatemia in the fed and fasted state (Figure 5A). Notably, the msKO and gKO models differ in that the latter is relatively protected against diet-induced obesity suggesting a role for other tissues such as adipose contributing to this lean phenotype. To further explore the impact of MondoA deficiency in the adipocyte, we generated human adipocytes from human adipocyte stem cells (hASCs) with CRISPR-generated knockdown (KD) of MondoA (Figure S6A). HPRT KD hASC derived adipocytes served as a control for the CRISPR process. Cultured MondoA KD hASC derived adipocytes demonstrated increased production of lactate compared to HPRT KD control adipocytes (Figure 5B). Additionally, using a 2-deoxyglucose assay, MondoA KD adipocytes demonstrated increased unstimulated and insulin-stimulated glucose uptake (Figure 5C), comparable to prior data in MondoA-deficient myotubes [2]. Increased glycolytic flux was further confirmed using a Seahorse assay for extracellular acidification rates (ECAR). MondoA KD adipocytes demonstrated increased maximal respiration, glycolytic capacity and unchanged reserve (Figure 5D). This effect was also confirmed by CRISPR editing downstream targets ARRDC4 and TXNIP KD in hASC derived adipocytes (Figure S6B-C) providing additional evidence that the increased glycolytic flux is related to the absence of the MondoA-Txnip/Arrdc4 inhibitory feedback on cellular glucose uptake.

**Fig. 5.**
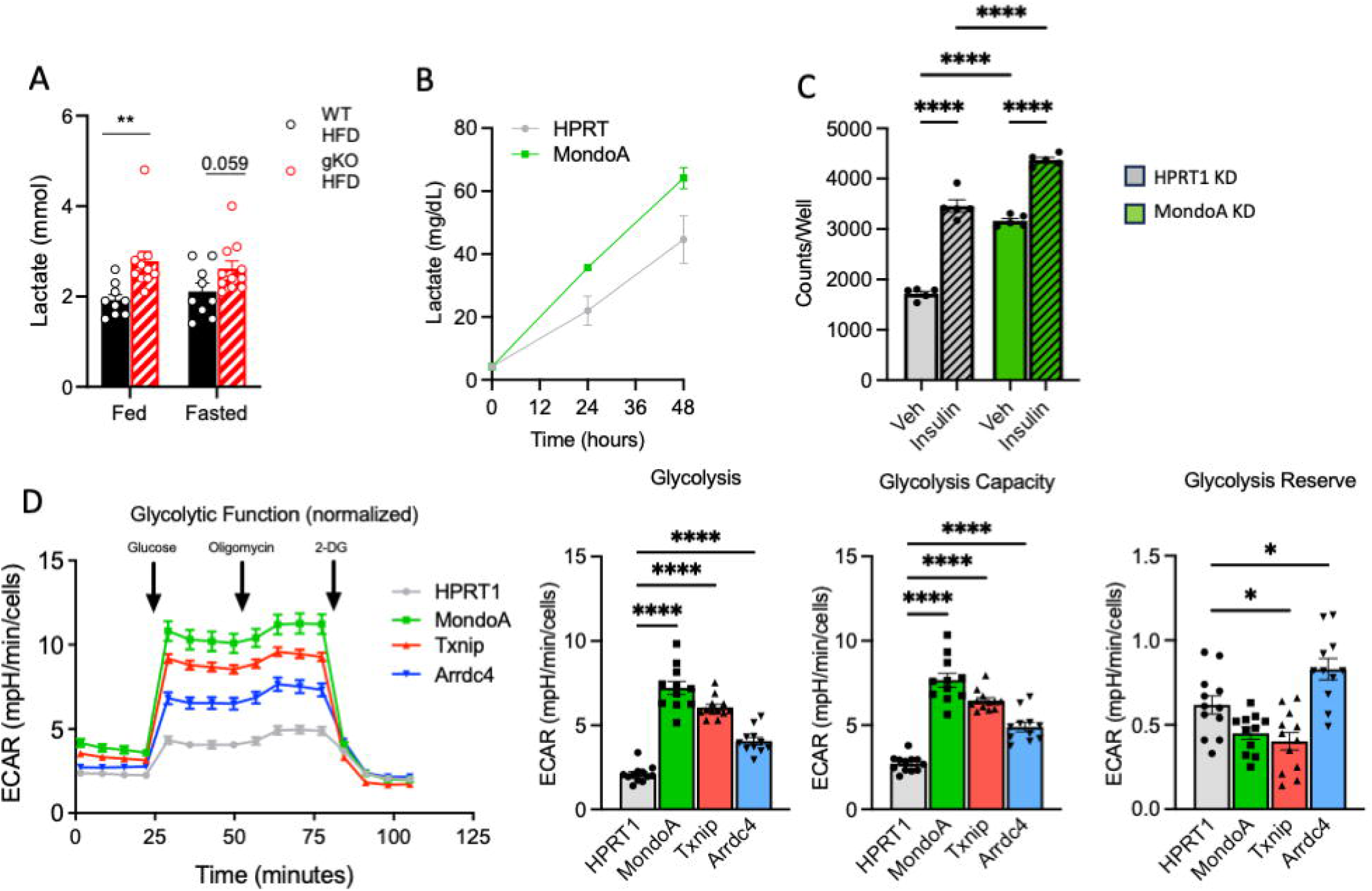
MondoA deficiency leads to increased energy expenditure which involves increased glycolytic flux. A) Whole blood lactate levels in HFD-fed WT and gKO male littermates (n=9-11C). B) Measurement of lactate generation from culture media of CRISPR-generated human iPSC-derived control (HPRT1) and MondoA KO adipocytes (hASCs). C) 2-deoxyglucose uptake assay with and without insulin in hASCs. D) Extracellular acidification rate (ECAR) in MondoA, Txnip and Arrdc4 KO hASCs and controls (HPRT1), representative from 3 independent experiments (n=10-11). Mean maximal respiration, glycolytic capacity and reserve ± SEM are quantified (right). CD, chow diet; HFD, high fat diet; KO, knockout; WT, wildtype. Data displayed as mean ± SEM. Student’s two-tailed t-test, 1-way, or 2-way ANOVA with Tukey’s comparison. P values * < 0.05, **<0.01, ****<0.001 displayed on graphs.

### MondoA deficiency in vitro results in altered stimulated lipolysis consistent with futile re-esterification

The hASC lines were next tested for capacity to lipolyze, synthesize, and oxidize triglyceride. Upon measuring isoproterenol-stimulated lipolysis, MondoA KD hASCs had significantly increased glycerol release and lower FFA release, compared to controls (Figure 6A). This decrease in isoproterenol-stimulated FFA release in MondoA KD cells was recapitulated across a wide range of isoproterenol doses, and MondoA KD cells also had lower maximal isoproterenol-stimulated FFA release (Figure 6B). We reasoned that this difference in glycerol:FFA ratio in MondoA KD cells could be a result of either increased fatty acid oxidation or increased fatty acid re-esterification. To understand which of these mechanisms may be affected in MondoA KD cells, we used OCR to assess oxidative metabolism of glucose, glutamine, and pyruvate (Figure 6C). Surprisingly, MondoA KD cells had decreased basal and maximal mitochondrial respiration, which was mirrored by *Txnip*, but not *Arrdc4*, deletion (Figure 6D). Fatty acid oxidation rates were then directly assessed using oxidation of ^3^H-palmitate to ^3^H-H_2_O (Figure 6E). Consistent with the OCR results, there was a small but significant decrease in fatty acid oxidation in MondoA and *Txnip* KD hASCs (Figure 6D). To further investigate fatty acid oxidation and re-esterification, stimulated lipolysis assays were repeated with the addition of either etomoxir to inhibit fatty acid oxidation or triacsin C to inhibit re-esterification (Figure 6F). While treatment with etomoxir did not alter levels of FFA, consistent with results from the palmitate oxidation assay, treatment with triacsin C resulted in normalization of FFA release in MondoA and *Txnip* KD cells with controls, indicating a reliance on re-esterification/lipolysis for the observed phenotype. To assess re-esterification and lipid cycling, palmitate and acetate uptake assays were conducted under isoproterenol-stimulated conditions. Increased ^14^C-palmitate incorporation into the total lipid pool in MondoA KD cells (Figure 6G, left) further supported a role for fatty acid re-esterification in this mechanism. Additionally, ^14^C-acetate incorporation into the cellular lipid pool was also higher in MondoA KD cells (Figure 6G, right) suggesting de novo lipogenesis may also be increased in these cells as part of a lipid futile cycle. Taken together, these results were consistent with fatty acid re-esterification with glycerol replenishment via biosynthesis driven by enhanced glucose uptake.

**Fig. 6.**
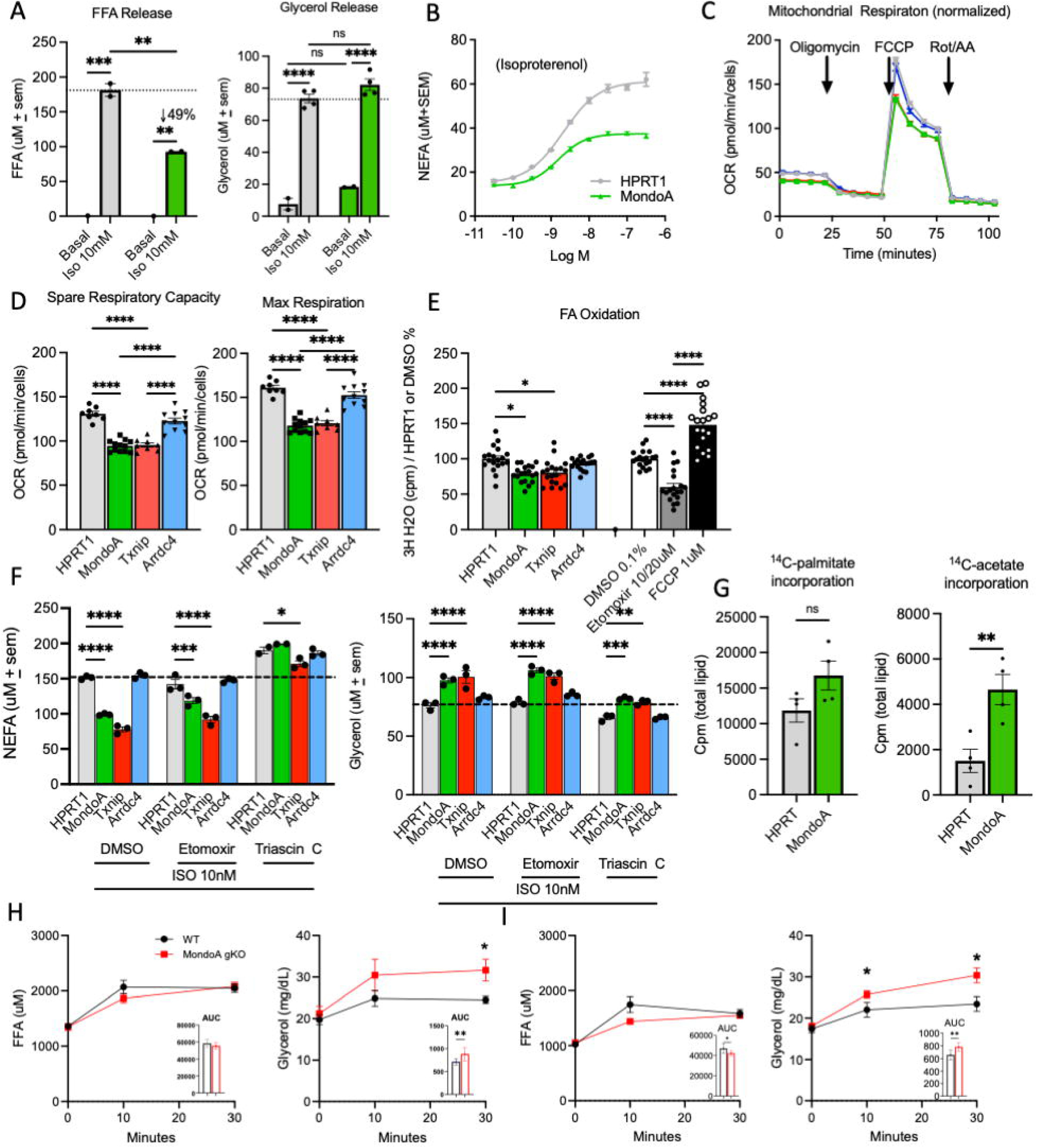
MondoA deficiency in mouse and human iPSC-derived adipocytes leads to isoproterenol-stimulated FFA release consistent with futile re-esterification. A) Basal and isoproterenol-stimulated glycerol and free fatty acid (FFA) release. B) Dose response curve for isoproterenol. C) Line graph represents oxygen consumption rates (OCR) in MondoA, Txnip or Arrdc4 KO APCs and controls (HPRT1), representative from 3 independent experiments (n=10-11). D) Bar graphs show the mean spare and maximal respiratory capacity ± SEM. E) In vitro fatty acid oxidation measured from 3H-palmitate in control, MondoA, Txnip and Arrdc4 KO APCs with positive and negative controls of Etomoxir and FCCP (n=9-13). F) Stimulated lipolysis assays for FFA and glycerol release in WT and KO APCs treated with Etomoxir and Triascin C. G) Incorporation of labelled substrate into the total lipid pool. Free fatty acid (FFA) and glycerol serum levels over time in ad lib H) chow-fed and I) HFD –fed WT and gKO male littermates following CL-316,243 stimulated lipolysis. Inset, area under the curve. The data represent mean ± SEM. All statistical significance determined by 1-way ANOVA with Tukey’s multiple-comparisons post hoc test (A, D, E, F) or 2-tailed student’s t-test (G, H,I). P values * < 0.05, ** < 0.01, *** <0.001, ****<0.0001, displayed on graphs.

Disproportionate glycerol:FFA ratio seen in the human-derived adipocytes has previously been shown to be an indicator of futile re-esterification - a futile cycling pathway [11]. To determine if this occurred *in vivo*, CD- and HFD-fed WT and gKO underwent a stimulated lipolysis assay using the selective beta-3 adrenergic receptor agonist CL-316,243. At basal conditions, there was no measured difference in unstimulated circulating free fatty acids (FFA) and glycerol, the breakdown products of lipolysis (Figure 6 H-I). However, upon stimulation, while both gKO and WT mice had a nearly 2-fold increase in circulating FFA, the gKO mice had a substantially larger molar increase in blood glycerol concentrations (recapitulating the *in vitro* observation).

### The protection against diet-induced obesity is not conferred by an adipocyte autonomous mechanism

Our collective results strongly implicated an adipocyte lipolysis/re-esterification mechanism for increased energy expenditure in the gKO mice in the context of HFD. We next sought to determine whether this phenotype could be replicated by *in vivo* adipocyte MondoA deficiency. To this end, an adipocyte-specific MondoA KO (aKO) mouse line was generated using an adiponectin promoter-driven Cre recombinase [12] (Figure S7A). MondoA deletion was confirmed in isolated adipocytes, with concomitant reduction in Arrdc4 but not *Txnip*, similar to gKO adipose tissue (Figure S7B). aKO and WT littermates were challenged with HFD feeding for 10 weeks, but no difference in weight gain was observed at standard room temperature (data not shown). In case the phenotype was subtle, the HFD challenge was repeated in the setting of thermoneutrality without any notable effect (Figure S7C). No differences in glucose or insulin handling were observed in the obese aKO mice compared to WT littermates (Figure S7D-E). These mice did not demonstrate increased lactatemia (Figure S7F). Finally, the aKO model underwent stimulated lipolysis testing, but did not demonstrate a difference to recapitulate the gKO mice (Figure S7G). These results indicate that MondoA deficiency in multiple tissues is required for the observed increased energy expenditure in the context of chronic caloric excess.

## Discussion

Regulating whole-body energy homeostasis requires tight coordination between nutrient sensing and substrate utilization, balancing energy needs and environmental inputs. Opposing metabolic pathways operating simultaneously across multiple organs create a futile cycle, leading to continued energy expenditure without obvious purpose (unless the goal is to produce heat). MondoA is a glucose-sensing transcription factor, inducing expression of *Txnip* and *Arrdc4* to decrease cellular glucose uptake, and shuttling glucose and fatty acids into glycogen and triglyceride storage depots, respectively [1, 2]. Since this role has been well-studied in muscle but not in other tissues, we examined the results of whole-body MondoA deficiency. The gKO mice exhibit increased energy expenditure in the context of chronic caloric excess, and our results are consistent with an energy-expending mechanism that involves futile cycling between adipose and other tissues (see Graphical Abstract).

MondoA deficiency results in loss of nutrient sensing and shuttling of nutrients into storage depots in the context of caloric excess by regulating expression of its target genes *Txnip* and *Arrdc4*. The MondoA/Txnip-Arrdc4 axis places a rapid positive-feedback brake on glucose uptake. Adipocytes and myocytes [2] display a robust increase in insulin-independent glucose uptake in the context of MondoA deficiency. At an organismal level, this results in improved insulin sensitivity and glucose tolerance in the gKO model, a phenotype shared by the msKO mice. Seahorse analyses also demonstrated that loss of MondoA/Txnip-Arrdc4 enhances glycolytic flux and high rates of glycolysis in muscle and adipose (and possibly other tissues) driving increased lactate production. Lactate is an important gluconeogenic substrate producing glucose for glycerol synthesis in adipocytes, a process akin to the Cori cycle in which exercise drives lactate-induced glucose production for muscle [25].

We also found that MondoA deficiency results in reduced lipid stores in both liver and adipose tissue. MondoA gKO mice exhibited reduced adiposity likely due to increased energy expenditure since caloric intake, fatty acid oxidation, adaptive thermogenesis, and physical activity were unchanged. The increased energy expenditure may be the result of futile cycling since we detected increased glycerol:FFA ratios in response to lipolytic stimuli *in vitro* and *in vivo*. Since liberated glycerol is released from the adipocyte at a faster rate than re-esterification can occur, for this cycle to be perpetuated, excess glucose is required to drive *de novo* glycerol production. At the same time, the increased lactate production from muscle and the glycerol secretion from adipose can drive *de novo* gluconeogenesis in the liver, feeding back into this futile cycle. We speculate that increased lipolysis and re-esterification may serves as an adaptive mechanism against hazardous lipid and lipid by-product accumulation.

Analyses of gene expression signatures across multiple tissues do not provide a “smoking gun” towards understanding how this futile cycle is established or maintained (other than the lack of Txnip and Arrdc4). It is worth speculating that many of the metabolic pathways may not be transcriptional in nature, but likely controlled by substrate flux and allosteric regulation across multiple tissues. In what appears to be an interesting observation of species differences, the MondoA-Arrdc4 interaction drives control in mice, while the MondoA-Txnip interactions appears more important in human-derived cells. Future studies using complex, multi-tracer labeling studies will need to be performed to demonstrate futile lipid cycling, but their practical application is limited [26].

Our experimental data in multiple tissues and model systems coalesce on a common conclusion: Glycerol-fatty acid cycling results in a state of increased energy expenditure and protection from diet-induced obesity. Critically, though there is a strong signature for adipose depot dysfunction, the observations in the global KO are not recapitulated in the adipose-specific KO mice. This implies a dependence on inter-organ crosstalk for initiating and sustaining this futile cycling. One form of inter-organ cross-talk involves the contribution of increased whole-body glycolysis to drive lactate which in turn can fuel hepatic gluconeogenesis to produce glucose for continued glycerol synthesis in adipocytes. The difference between the culture adipocytes and aKO mice may be related to developmental timing of MondoA deletion or an undefined signaling pathway.

In addition to a critical glucose sensing role, MondoA modulates the inflammatory/pro-oxidant response via *Txnip* inhibition of thioredoxin and interaction with the NLRP3 inflammasome. These actions implicate MondoA in a multifaceted role for progressive inflammatory diseases, including insulin resistance and diabetic nephropathy [1, 2, 27], ischemic cardiac disease [28], neurodegenerative conditions [29], cancer [30, 31], and cellular aging [32]. It is unclear how these diverse disease states relate to our observations of critical multi-organ crosstalk for nutrient sensing and regulation, though intriguing to speculate about metabolic underpinnings. Importantly, MondoA inhibition has a generally favorable effect in the setting of nutrient excess. Our lab has previously demonstrated that a small-molecular, SBI-993, can inhibit MondoA action; given the results herein and the success of novel systemic pharmacologic agents like the SGLT2 inhibitors and GLP-1 agonists, targeting MondoA may have promise in metabolic and other disease states.

### Conclusions

Global MondoA deficiency results in resistance to diet-induced obesity due to increased energy expenditure and cold tolerance, which we hypothesize is due to futile glycerol-FFA cycling. Critical findings from this conceptual model can be replicated in human adipocytes. Differences in mouse and human MondoA target preference, and absence of this phenotype in adipose- or skeletal muscle-specific KO mice suggest a profound organismal level homoeostasis which requires further study. MondoA and its targets are a viable and intriguing therapeutic checkpoint for energy balance in human disease.

## Acknowledgements

We thank Teresa C. Leone, Ling Lai, and Russell Callaway for technical assistance. Indirect calorimetry was performed by the Rodent Metabolic Phenotyping Core at UPenn supported in part by NIH grant S10-OD025098 and the Cox Institute. Histology was prepared by the Molecular Pathology and Imaging Core at the Penn Center for Molecular Studies in Digestive and Liver Diseases (RRID: SCR_022420) supported in part by NIH grant P30DK050306. Adipose histology was performed by Sarah Traynor and David Merrick with assistance from Lan Cheng and the Penn IDOM histology core. Biorender.com was used for creation of figure schematics.

## CRediT authorship contribution statement

JHB: Conceptualization, Formal Analysis, Investigation, Writing – original draft; ANL: Investigation, Formal Analysis, Writing – original draft; LCJ: Investigation, Formal Analysis, Writing – original draft; RT: Formal Analysis, Investigation, Writing – review & editing; BA: Conceptualization, Investigation; XY: Investigation; TS: Investigation, Methodology, Writing – review & editing; KB: Formal Analysis, Visualization; JP: Investigation; JZ: Investigation; PMT: Conceptualization, Writing – review & editing; BNF: Writing – review & editing; GT: Conceptualization, Validation, Writing – review & editing; DPK: Conceptualization, Funding acquisition, Validation, Writing – review & editing

## Data Sharing

All data have been deposited in NCBI’s Gene Expression Omnibus Series accession number GSE303998. The following link has been created to allow review prior to publication: https://www.ncbi.nlm.nih.gov/geo/query/acc.cgi?acc=GSE303998. Please use the following secure token to access the site: mjelcekqtfuzrkt.

## Footnotes

1 Non-standard abbreviations: BAT, brown adipose tissue; CD, chow diet; ECAR, Extra Cellular Acidification Rate; FFA, free fatty acid; hASC, human-derived adipocytes; GTT, glucose tolerance test; HFD, high fat diet; ITT, insulin tolerance test; KEGG, Kyoto Encyclopedia of Genes and Genomes; KD, knock-down; KO, knockout (generalized, gKO, muscle-specific, msKO, or adipose-specific aKO); OCR, Oxygen Consumption Rate; PCA, principal component analysis; PTT, pyruvate tolerance test; WAT, white adipose tissue (iWAT, inguinal, or eWAT, epidydimal); WT, wildtype.

## References

[1] Ahn, B., Wan, S., Jaiswal, N., Vega, R.B., Ayer, D.E., Titchenell, P.M., et al., 2019. MondoA drives muscle lipid accumulation and insulin resistance. JCI Insight 5(15).

[2] Ahn, B., Soundarapandian, M.M., Sessions, H., Peddibhotla, S., Roth, G.P., Li, J.L., et al., 2016. MondoA coordinately regulates skeletal myocyte lipid homeostasis and insulin signaling. J Clin Invest 126(9):3567–3579.

[3] Imamura, M., Chang, B.H., Kohjima, M., Li, M., Hwang, B., Taegtmeyer, H., et al., 2014. MondoA deficiency enhances sprint performance in mice. Biochem J 464(1):35–48.

[4] Chen, J.L., Merl, D., Peterson, C.W., Wu, J., Liu, P.Y., Yin, H., et al., 2010. Lactic acidosis triggers starvation response with paradoxical induction of TXNIP through MondoA. PLoS Genet 6(9):e1001093.

[5] Wilde, B.R., Ye, Z., Lim, T.Y., Ayer, D.E., 2019. Cellular acidosis triggers human MondoA transcriptional activity by driving mitochondrial ATP production. Elife 8.

[6] Sharma, A.K., Khandelwal, R., Wolfrum, C., 2024. Futile cycles: Emerging utility from apparent futility. Cell Metab 36(6):1184–1203.

[7] Vargas-Castillo, A., Sun, Y., Smythers, A.L., Grauvogel, L., Dumesic, P.A., Emont, M.P., et al., 2024. Development of a functional beige fat cell line uncovers independent subclasses of cells expressing UCP1 and the futile creatine cycle. Cell Metab 36(9):2146–2155 e2145.

[8] Kazak, L., Chouchani, E.T., Jedrychowski, M.P., Erickson, B.K., Shinoda, K., Cohen, P., et al., 2015. A creatine-driven substrate cycle enhances energy expenditure and thermogenesis in beige fat. Cell 163(3):643–655.

[9] Oeckl, J., Janovska, P., Adamcova, K., Bardova, K., Brunner, S., Dieckmann, S., et al., 2022. Loss of UCP1 function augments recruitment of futile lipid cycling for thermogenesis in murine brown fat. Mol Metab 61:101499.

[10] Chitraju, C., Mejhert, N., Haas, J.T., Diaz-Ramirez, L.G., Grueter, C.A., Imbriglio, J.E., et al., 2017. Triglyceride Synthesis by DGAT1 Protects Adipocytes from Lipid-Induced ER Stress during Lipolysis. Cell Metab 26(2):407–418 e403.

[11] Rotondo, F., Ho-Palma, A.C., Remesar, X., Fernandez-Lopez, J.A., Romero, M.D.M., Alemany, M., 2017. Glycerol is synthesized and secreted by adipocytes to dispose of excess glucose, via glycerogenesis and increased acyl-glycerol turnover. Sci Rep 7(1):8983.

[12] Eguchi, J., Wang, X., Yu, S., Kershaw, E.E., Chiu, P.C., Dushay, J., et al., 2011. Transcriptional control of adipose lipid handling by IRF4. Cell Metab 13(3):249–259.

[13] Mina, A.I., LeClair, R.A., LeClair, K.B., Cohen, D.E., Lantier, L., Banks, A.S., 2018. CalR: A Web-Based Analysis Tool for Indirect Calorimetry Experiments. Cell Metab 28(4):656–666 e651.

[14] Weir, J.B., 1949. New methods for calculating metabolic rate with special reference to protein metabolism. J Physiol 109(1-2):1–9.

[15] Merrick, D., Sakers, A., Irgebay, Z., Okada, C., Calvert, C., Morley, M.P., et al., 2019. Identification of a mesenchymal progenitor cell hierarchy in adipose tissue. Science 364(6438).

[16] Sakamoto, T., Batmanov, K., Wan, S., Guo, Y., Lai, L., Vega, R.B., et al., 2022. The nuclear receptor ERR cooperates with the cardiogenic factor GATA4 to orchestrate cardiomyocyte maturation. Nat Commun 13(1):1991.

[17] Yamamoto, T., Maurya, S.K., Pruzinsky, E., Batmanov, K., Xiao, Y., Sulon, S.M., et al., 2023. RIP140 deficiency enhances cardiac fuel metabolism and protects mice from heart failure. J Clin Invest 133(9).

[18] Patro, R., Duggal, G., Love, M.I., Irizarry, R.A., Kingsford, C., 2017. Salmon provides fast and bias-aware quantification of transcript expression. Nat Methods 14(4):417–419.

[19] Love, M.I., Huber, W., Anders, S., 2014. Moderated estimation of fold change and dispersion for RNA-seq data with DESeq2. Genome Biol 15(12):550.

[20] Kolberg, L., Raudvere, U., Kuzmin, I., Adler, P., Vilo, J., Peterson, H., 2023. g:Profiler-interoperable web service for functional enrichment analysis and gene identifier mapping (2023 update). Nucleic Acids Res 51(W1):W207–W212.

[21] Chen, Y., Liu, L., Calhoun, R., Cheng, L., Merrick, D., Steger, D.J., et al., 2025. Transcriptional regulation of adipocyte lipolysis by IRF2BP2. Sci Adv 11(1):eads5963.

[22] Sakamoto, T., Nitta, T., Maruno, K., Yeh, Y.S., Kuwata, H., Tomita, K., et al., 2016. Macrophage infiltration into obese adipose tissues suppresses the induction of UCP1 level in mice. Am J Physiol Endocrinol Metab 310(8):E676–E687.

[23] Hotamisligil, G.S., 2006. Inflammation and metabolic disorders. Nature 444(7121):860–867.

[24] Mittenbuhler, M.J., Jedrychowski, M.P., Van Vranken, J.G., Sprenger, H.G., Wilensky, S., Dumesic, P.A., et al., 2023. Isolation of extracellular fluids reveals novel secreted bioactive proteins from muscle and fat tissues. Cell Metab 35(3):535–549 e537.

[25] Soeters, P.B., Shenkin, A., Sobotka, L., Soeters, M.R., de Leeuw, P.W., Wolfe, R.R., 2021. The anabolic role of the Warburg, Cori-cycle and Crabtree effects in health and disease. Clin Nutr 40(5):2988–2998.

[26] Wunderling, K., Zurkovic, J., Zink, F., Kuerschner, L., Thiele, C., 2023. Triglyceride cycling enables modification of stored fatty acids. Nat Metab 5(4):699–709.

[27] Tan, C.Y., Weier, Q., Zhang, Y., Cox, A.J., Kelly, D.J., Langham, R.G., 2015. Thioredoxin-interacting protein: a potential therapeutic target for treatment of progressive fibrosis in diabetic nephropathy. Nephron 129(2):109–127.

[28] Domingues, A., Jolibois, J., Marquet de Rouge, P., Nivet-Antoine, V., 2021. The Emerging Role of TXNIP in Ischemic and Cardiovascular Diseases; A Novel Marker and Therapeutic Target. Int J Mol Sci 22(4).

[29] Nasoohi, S., Ismael, S., Ishrat, T., 2018. Thioredoxin-Interacting Protein (TXNIP) in Cerebrovascular and Neurodegenerative Diseases: Regulation and Implication. Mol Neurobiol 55(10):7900–7920.

[30] Carroll, P.A., Diolaiti, D., McFerrin, L., Gu, H., Djukovic, D., Du, J., et al., 2015. Deregulated Myc requires MondoA/Mlx for metabolic reprogramming and tumorigenesis. Cancer Cell 27(2):271–285.

[31] Sipol, A., Hameister, E., Xue, B., Hofstetter, J., Barenboim, M., Ollinger, R., et al., 2022. MondoA drives malignancy in B-ALL through enhanced adaptation to metabolic stress. Blood 139(8):1184–1197.

[32] Yamamoto-Imoto, H., Minami, S., Shioda, T., Yamashita, Y., Sakai, S., Maeda, S., et al., 2022. Age-associated decline of MondoA drives cellular senescence through impaired autophagy and mitochondrial homeostasis. Cell Rep 38(9):110444.

