## Supplemental figures for "Whole body MondoA deletion protects against diet-induced obesity through uncontrolled multi-organ substrate utilization and futile cycling"

**Supplemental Fig 1. Generation of a whole-body MondoA knockout shows tissue specific gene changes.** A) Schema for generation of a generalized MondoA knockout mouse. B) QT-PCR in tissues from chow and HFD-fed mice for MondoA, and direct targets Txnip and Arrdc4 (n=6-10). HFD, high fat diet; KO, knockout; WT, wildtype. P values * < 0.05, *** <0.001, ****<0.0001 displayed on graphs. The data represent mean ± SEM. All statistical significance determined by two-way analysis of variance (ANOVA) with Tukey's multiple-comparisons post hoc test.

---

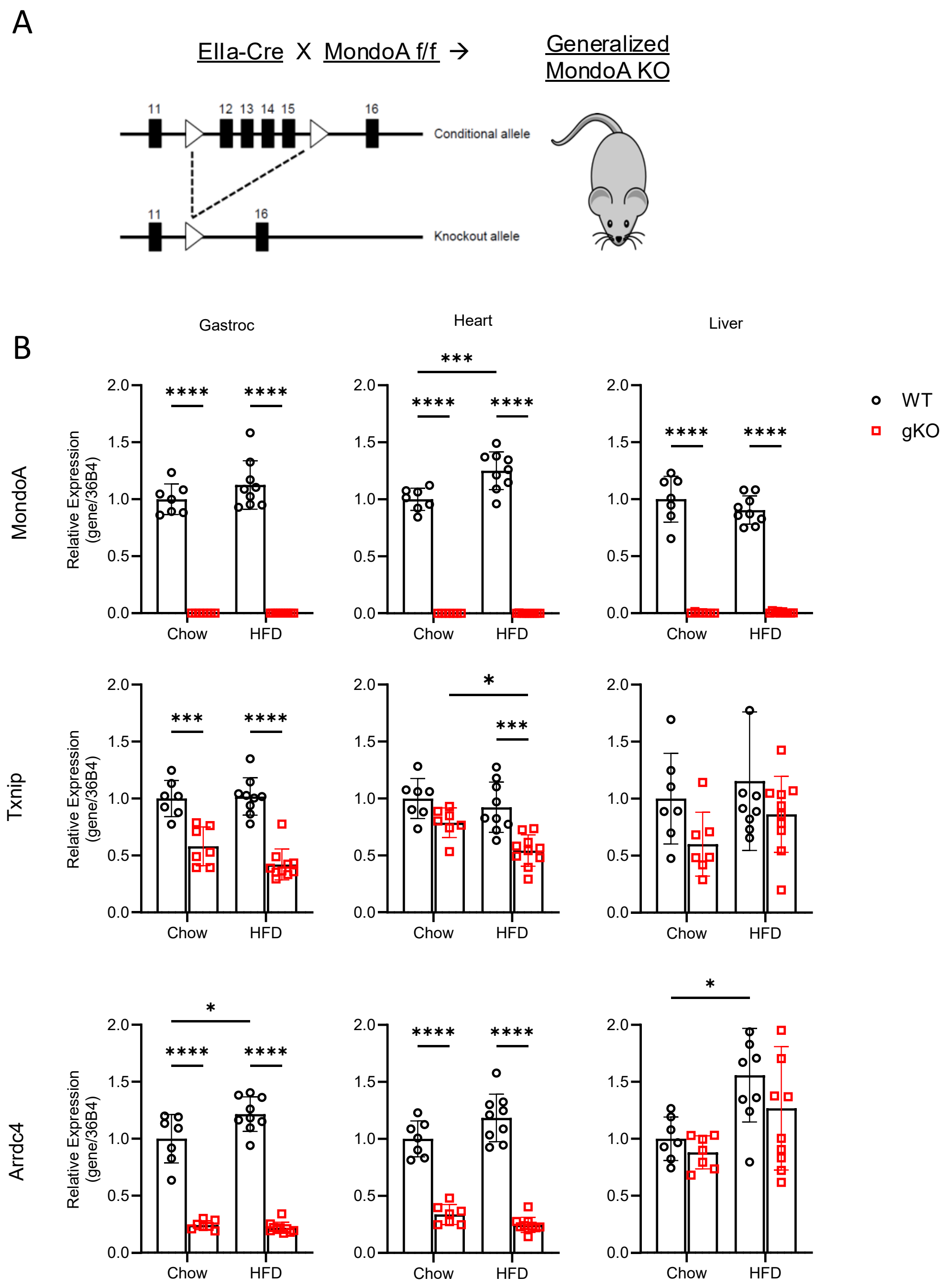

**Supplemental Fig 2. HFD-fed MondoA gKO mice have reduced hepatosteatosis.** Representative gross photographs and microscopy of chow and HFD-fed WT and gKO liver. CD, chow diet; H&E, hematoxylin and eosin staining; HFD, high fat diet; KO, knockout; ORO, oil-red O staining; WT, wildtype.

---

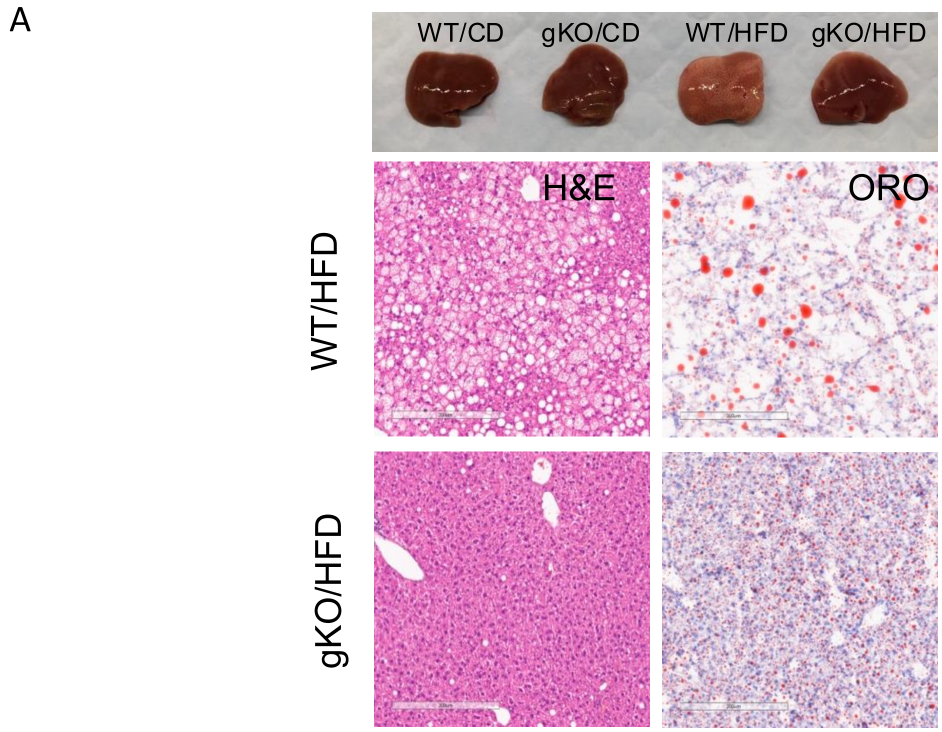


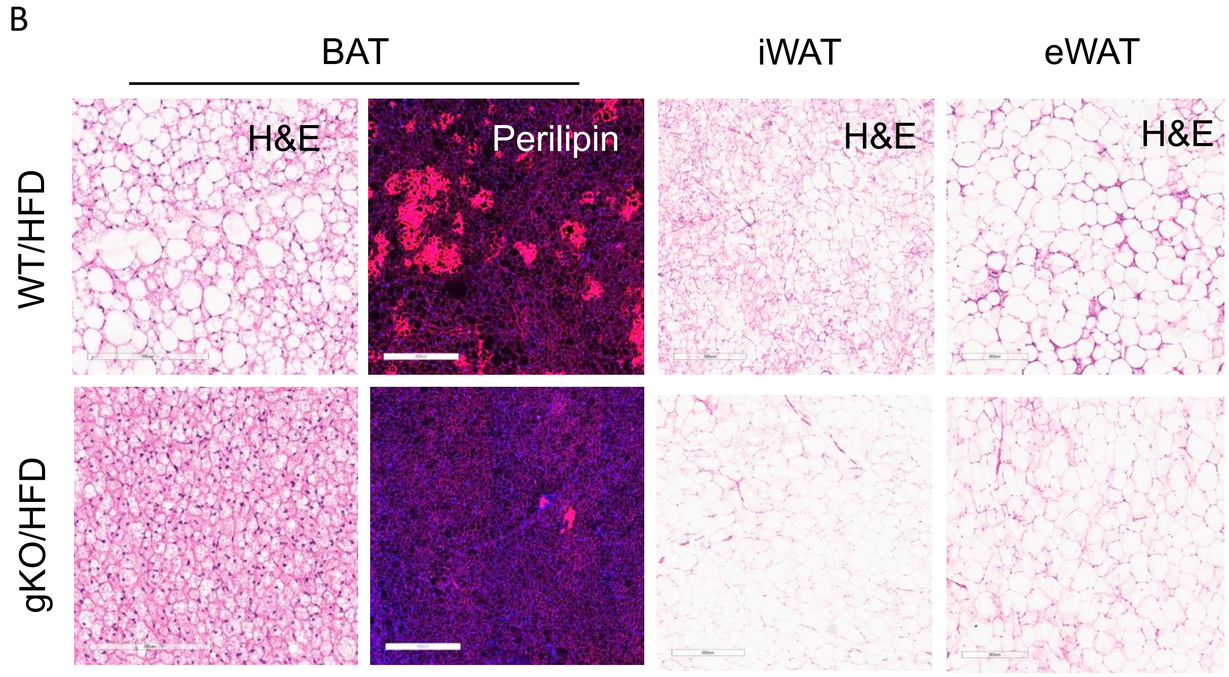

**Supplemental Fig 3. Whole-body MondoA deficiency shows adipose depot-specific gene changes.** QT-PCR in brown (BAT), inguinal white (iWAT) and epididymal white (eWAT) adipose tissues from chow and HFD-fed mice for MondoA, and direct targets Txnip and Arrdc4 (n=6-10). HFD, high fat diet; KO, knockout; WT, wildtype. Data displayed as mean ± SEM. Two-way analysis of variance (ANOVA) followed by multiple-comparisons test. P values * < 0.05, **<0.01. *** <0.001, ****<0.0001 displayed on graphs.

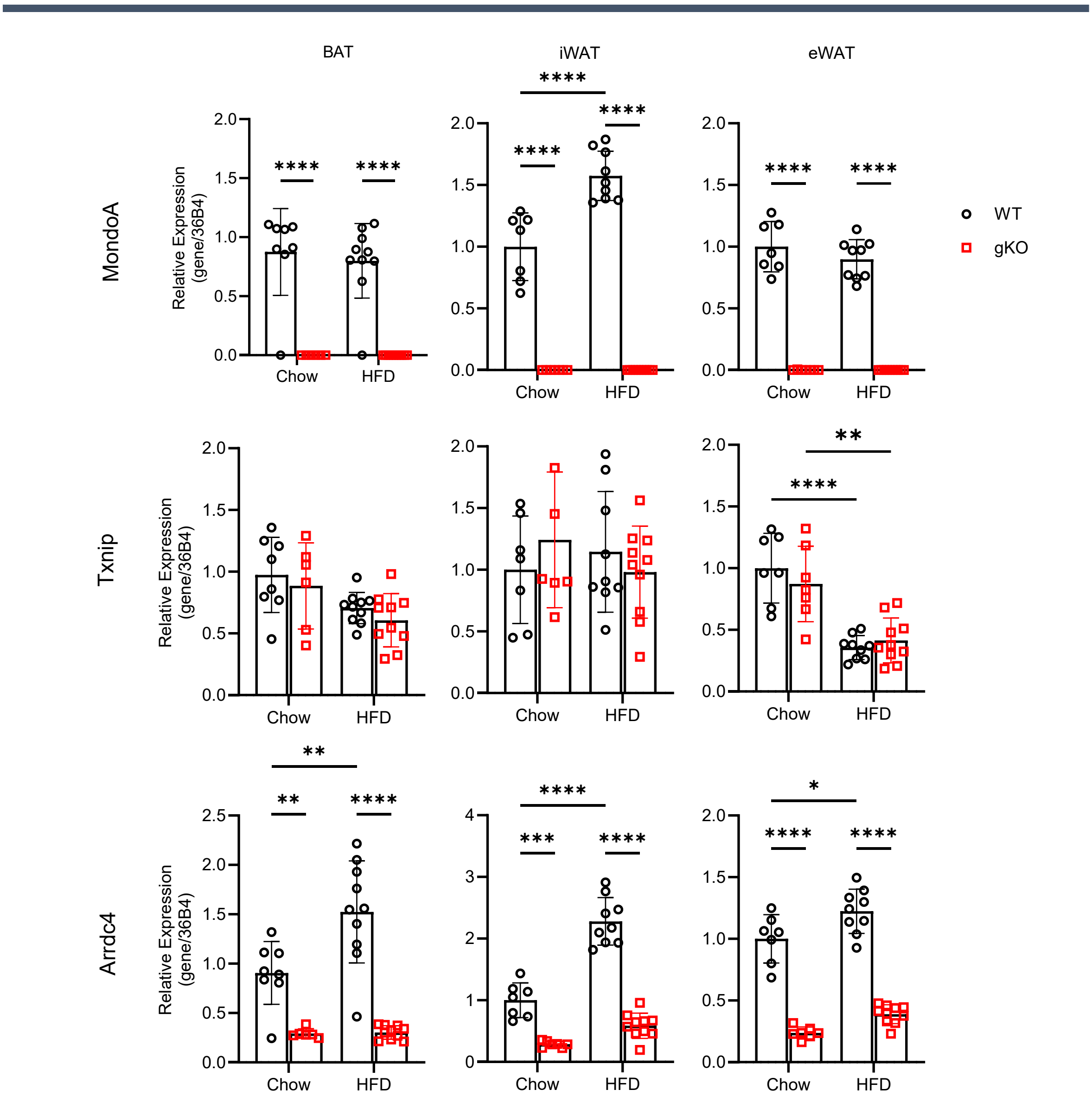

**Supplemental Fig. 4. Transcriptional analysis of MondoA gKO effects in BAT and iWAT**. Principal component analysis (PCA) plot (top) and heat map for (bottom) differentially expressed genes from RNAseq of A) BAT (n=4) and B) iWAT (N=6) in chow and HFD-fed WT and MondoA littermates. C) KEGG analysis and D) RPKM of differentially express genes from BAT. CD, chow diet; HFD, high fat diet; KO, knockout; WT, wildtype. Data displayed as mean ± SEM. Student's t-test or two-way analysis of variance (ANOVA) followed by multiple-comparisons test. P values * < 0.05, **< 0.01, *** <0.001, ****<0.0001 displayed on graphs.

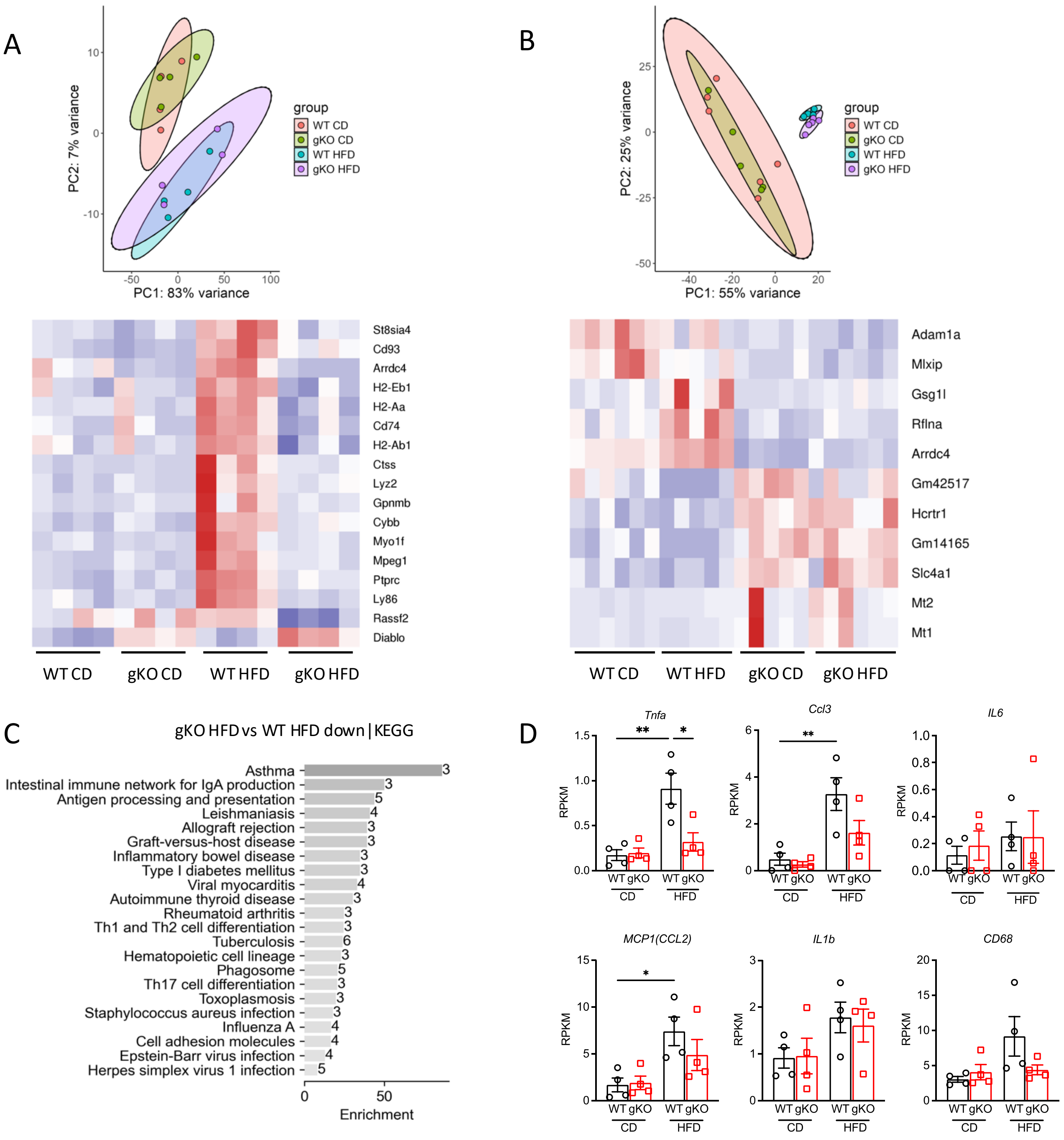

**Supplemental Fig. 5. Transcriptional analysis of non-classical adaptive thermogenesis in skeletal muscl**e and iWAT. A) QT-PCR targets for non-shivering thermogenesis from gastrocnemius isolated from chow-fed WT and MondoA gKO male littermates (n=5-6) maintained at thermoneutrality for 14 days prior to acute cold exposure. B) QT-PCR targets of creatine futile cycling in iWAT from chow and HFD-fed WT and gKO mice (same cohort as Fig. 5C-D and Sup. Fig 3B). gKO, knockout; TN, thermoneutrality; WT, wildtype. Data displayed as mean ± SEM. Student's t-test of 2-way ANOVA. P values * < 0.05, *** <0.001 displayed on graphs.

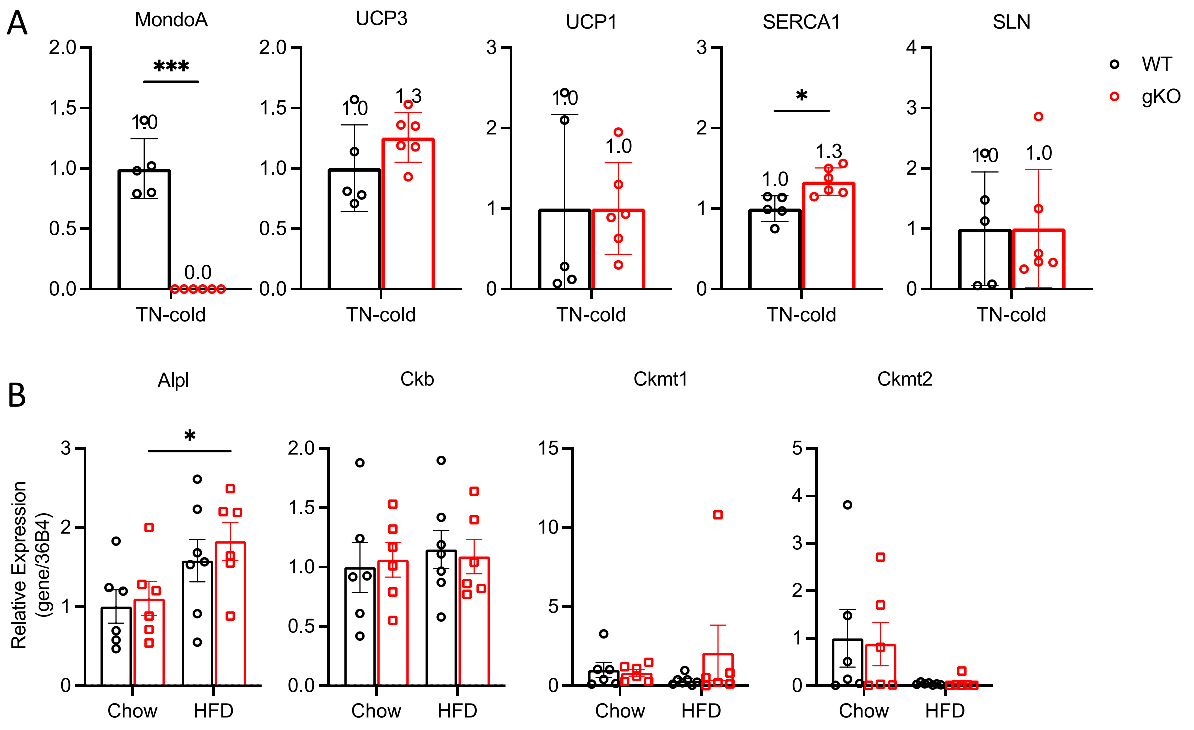

**Supplemental Fig. 6. Validation of CRISPR knockdown in human stem cell-derived adipocytes.** Western blot and quantification of A) MondoA and B) Txnip CRISPR KD in hASCs. C) QT-PCR of *Arrdc4* in hASCs (given the lack of a satisfactory ARRDC4 antibody). Data displayed as mean ± SEM. 1-way ANOVA. P values * < 0.05, **< 0.01, *** <0.001 displayed on graphs.

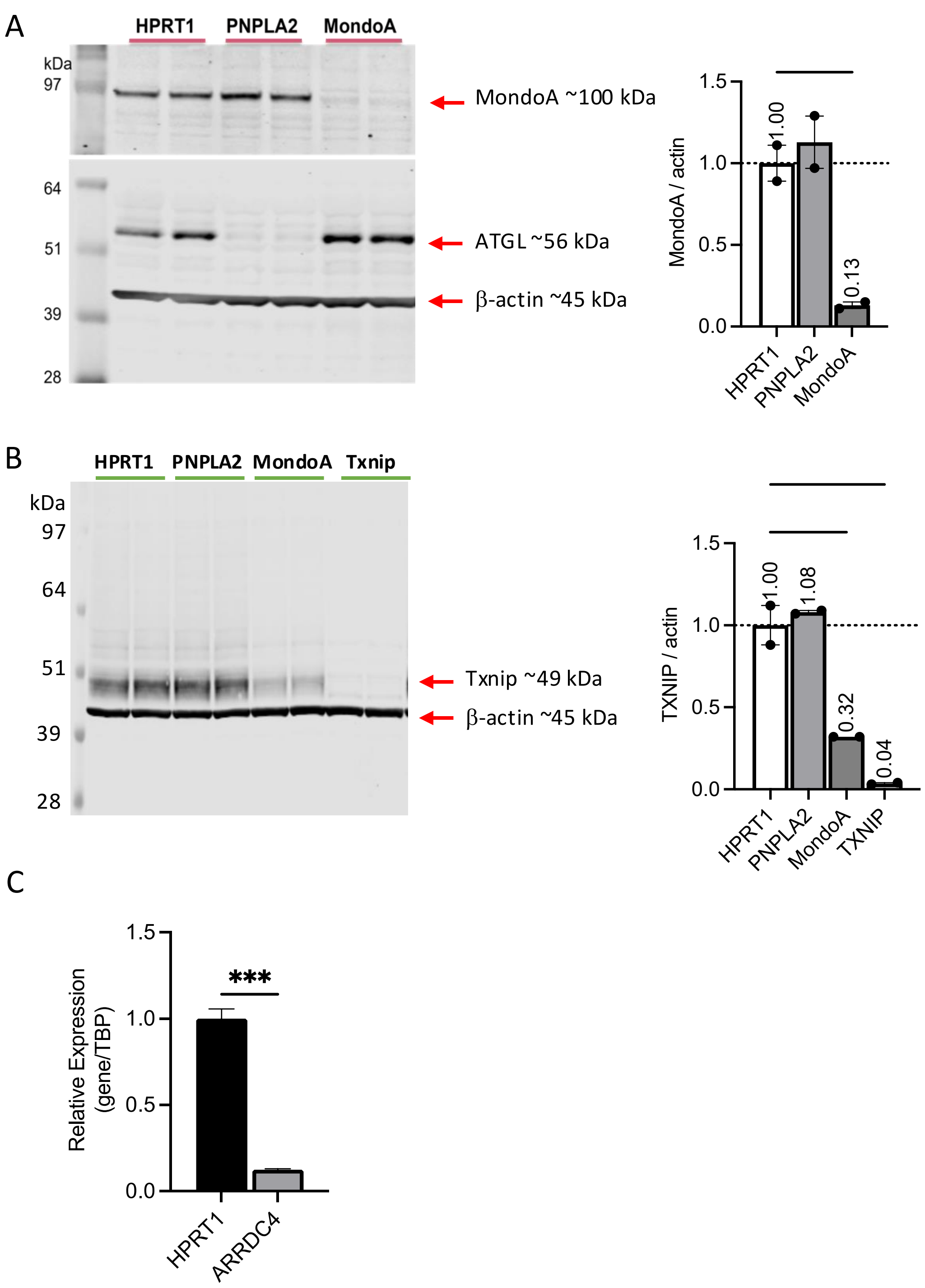

**Supplemental Fig. 7. Adipose-specific MondoA KO mouse does not have lean phenotype.** A) Schema for generation of an adipose-specific MondoA knockout (aKO) mouse. B) QT-PCR for cell isolation quality control, MondoA, and direct targets Txnip and Arrdc4 from adipocytes and stromal vascular cells isolated from HFD-fed WT and aKO mice (n=3). C) Body weight trends for WT and aKO MondoA mice on 60% kcal HFD over ten weeks at thermoneutrality (n=***). D) Glucose (left) and insulin (right) tolerance tests and area under the curve (inset). E) Fasting insulin levels. F) Random circulating lactate levels. G) Free fatty acid (FFA) and glycerol serum levels over time in ad lib HFD –fed WT and aKO male littermates following CL-316,243 stimulated lipolysis. Inset, area under the curve. HFD, high fat diet; KO, knockout; SVF, stromal vascular fraction; WT, wildtype. Data displayed as mean ± SEM. One-way analysis of variance (ANOVA) followed by multiple-comparisons test. P values displayed on graphs.

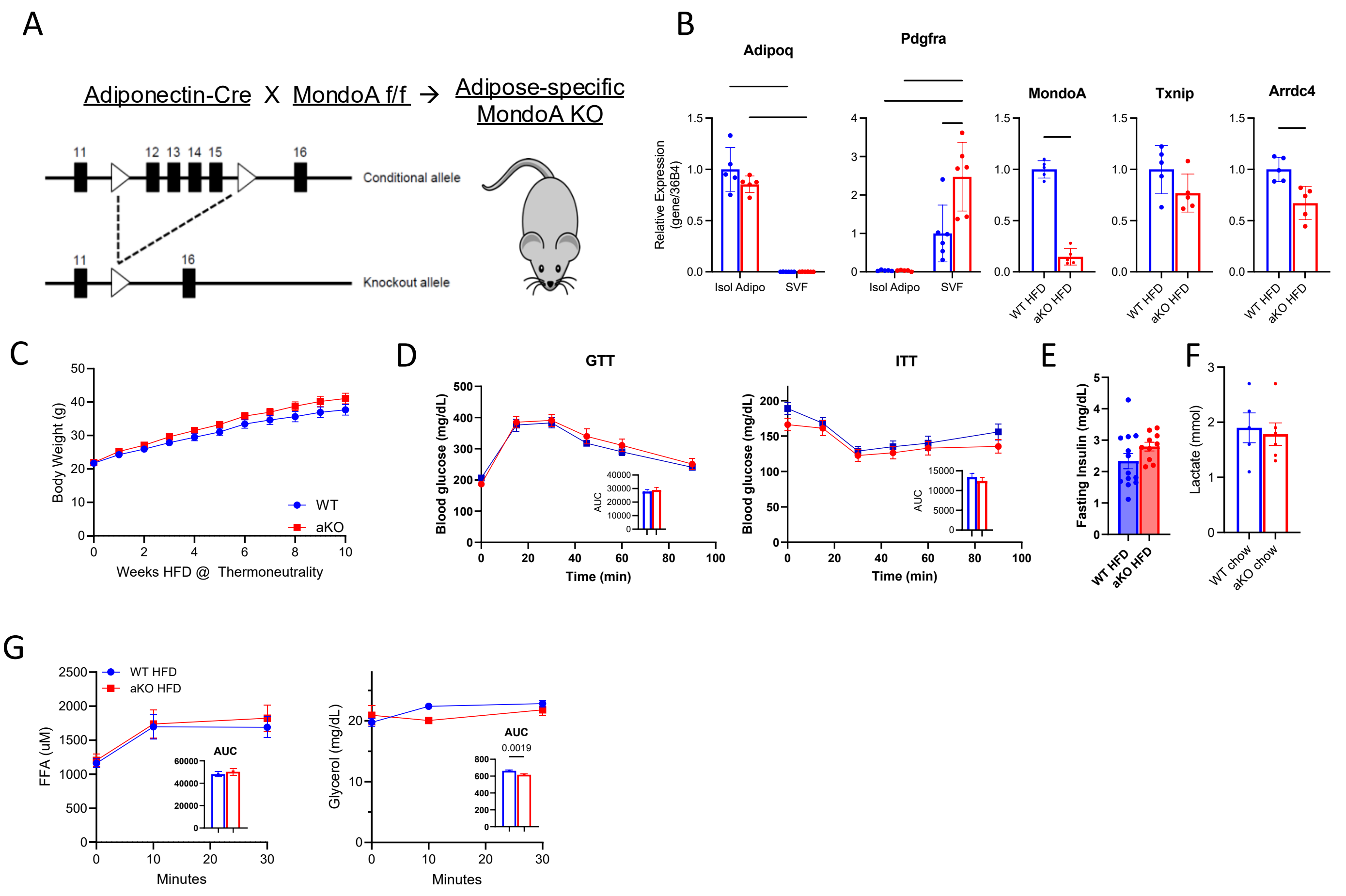

### Supp. Table 1: QT-PCR primers

| Gene | Forward | Reverse |
|---|---|---|
| Acadm | ATGACGGAGCAGCCAATGAT | TAATGGCCGCCACATCAGAG |
| Acsl1 | CGCCCATATGTTTGAGACCG | GTCGTCCATAAGCAGCCTGA |
| Adipoq | CTCCTGGAGAGAAGGGAGAGA | ACACATAAGCGGCTTCTCCAG |
| Arrdc4 | CAGCCTCCTCAGAAGTGGAAT | TCAGACGGAAGCTGAAAGCG |
| Atgl | GGATGGCGGCATTTCAGACA | CAAAGGGTTGGGTTGGTTCAG |
| CD36/Fat | CGCACATTGAGATTCTTTTCC | TCCTTTAAGGTCGATTTCAGATC |
| Cidea | TGACATTCATGGGATTGCAGAC | GGCCAGTTGTGATGACTAAGAC |
| Dgat1 | GCGACGGCTACTGGGATCTG | TGCATTACTCAGGATCAGCATCA |
| Dio2 | CAGTGTGGTGCACGTCTCCAATC | TGAACCAAAGTTGACCACCAG |
| Elovl3 | TTCTCACGCGGGTTAAAAATGG | GAGCAACAGATAGACGACCAC |
| Elovl6 | CAGTCAGTTTGTGACCAGAGTTTTTAC | CAGGAAGATCAGTTTCTGTTTCCTCAG |
| Fabp3 | AAGTGGAACGGGCAGGAGA | GAGGAGCGGGCGGTCAG |
| Fasn | CCAAGCAGGCACACACAATG | GATGCCTCTGAACCACTCACAC |
| Hadha | GAAAGCCAAGCCCAAAGA | TCAGGAGGGCTCAAAGAATAA |
| Hsl | CCAGCCTGAGGGCTTACTG | CTCCATTGACTGTGACATCTCG |
| Mlxip (MondoA) | TGCTACCTGCCACAGGAGTC | GACTCAAACAGTGGCTTGATGA |
| Pdgfra | ATCGTGGCTGAAGGACAACTT | TCCTTAGCCCGGATCAGCTT |
| Pdk4 | CCGCTGTCCATGAAGCA | GCAGAAAAGCAAAGGACGTT |
| Prdm16 | CAGCACGGTGAAGCCATTC | GCGTGCATCCGCTTGTG |
| Serca1 | TGTTTGTCCTATTTCGGGGTG | AATCCGCACAAGCAGGTCTTC |
| Sln | ACTGAGGTCCTTGGTAGCCT | AAGGACTTGTGATTGCACACC |
| Txnip | GTCTCAGCAGTGCAAACAGACTT | GCTCGAAGCCGAACTTGTACTC |
| Ucp1 | CGACTCAGTCCAAGAGTACTTCTCTTC | GCCGGCTGAGATCTTGTTTC |
| Ucp3 | TGCTGAGATGGTGACCTACGA | CCAAAGGCAGAGACAAAGTGA |
| | ThermoFisher Scientific ID# | |
| Arrdc4 | Hs00411771_m1 | |
| TBP | Hs00427620_m1 | |

**Supp. Table 2: sgRNA sequences for CRISPR/Alt-R S.p. Cas9 Nuclease V3 knockouts**

| Target | Guide sequence (Alt-R CRISPR-Cas9 sgRNA) | |
|---|---|---|
| **HPRT1 (Non-targeting control)** | 5'- AAT TAT GGG GAT TAC TAG GA -3' | http://www.ensembl.org/id/ENSG00000165704 |
| **MLXIP** | 5'- GTG TCC ATG AAG TCC AGG TT -3' | http://www.ensembl.org/id/ENSG00000175727 |
| **TXNIP** | 5'- GTT GAA TAT TCC TTA CTG GT -3' | http://www.ensembl.org/id/ENSG00000265972 |
| **ARRDC4** | 5'- TCC CGC TTT ACA CTC TGA TC -3' | http://www.ensembl.org/id/ENSG00000140450 |

**Supp. Table 3: Genomic PCR and Sequencing Primer sequences**

| Gene | Primer | Sequence 5'→3' |
|---|---|---|
| **HPRT1** | forward | TGA ACC CAG GAG GCA GAG C |
| | reverse | GGT CCT TTT CAG CAG CAA GCT G |
| | sequencing | TGC TCA CCT CTC CCA CAC CC |
| **MLXIP** | forward | GTG CGG AAT GGC CAC AAA CC |
| | reverse | GGA GAA CTG CTT ACA GTC ACG GC |
| | sequencing | AGC CAG CTC CTT GGA AGA CTC |
| **TXNIP** | forward | GGG TGT CTG TCT CTG CTC GAA |
| | reverse | CTG ATC TGC TGC CAA TTA CCA GGG |
| | sequencing | GGG ACT GGG GTA GAG GAA GTG TC |
| **ARRDC4** | forward | TGT GGT TAT CCT TCT CTG CAG T |
| | reverse | AAA GAA GCC CTG AAC AGC AG |
| | sequencing | AGC CAT CAG AAA GTG ACA GGA |

**Supp. Table 4: Antibodies**

| Antibody | Product number | Concentration |
|---|---|---|
| **MondoA** | Bethyl Labs A303-195AM | 1:1000 |
| **Txnip** | Cell Signaling #14715 | 1:1000 |
| **ATGL** | Cell Signaling #2138 | 1:1000 |
| **B-actin** | Sigma S5441 | 1:2000 |
| **Perilipin** | Cell Signaling #3470 | 1:500 |
